# Prolonged signaling of backbone-modified glucagon-like peptide-1 analogues with diverse receptor trafficking

**DOI:** 10.1101/2024.04.17.589632

**Authors:** Brian P. Cary, Marlies V. Hager, Rylie K. Morris, Matthew J. Belousoff, Patrick M. Sexton, Denise Wootten, Samuel H. Gellman

## Abstract

Signal duration and subcellular location are emerging as important facets of G protein-coupled receptor (GPCR) function. The glucagon-like peptide-1 receptor (GLP-1R), a clinically relevant class B1 GPCR, stimulates production of the second messenger cAMP upon activation by the native hormone, GLP-1. cAMP production continues after the hormone-receptor complex has been internalized via endocytosis. Here, we report GLP-1 analogues that induce prolonged signaling relative to GLP-1. A single β-amino acid substitution at position 18, with the residue derived from (*S*,*S*)-*trans*-2-aminocyclopentanecarboxylic acid (ACPC), enhances signaling duration with retention of receptor endocytosis. Pairing ACPC at position 18 with a second substitution, α-aminoisobutyric acid (Aib) at position 16, abrogates endocytosis, but prolonged signaling is maintained. Prolonged signaling is sensitive to the structure of the β residue at position 18. Cryo-electron microscopy (cryo-EM) structures of two GLP-1 analogues bound to the GLP- 1R:Gs complex suggest substantial alterations to bound peptide structure and dynamics compared to the GLP-1:GLP-1R complex. These structural findings strengthen an emerging view that agonist dynamics in the receptor-bound state influence signaling profile. Our results advance understanding of the structural underpinnings of receptor activation and introduce new tools for exploring the impact of spatiotemporal signaling profiles following GLP-1R activation.

## Introduction

The timeframe and site(s) of signaling mediated by G protein-coupled receptors (GPCRs) are increasingly recognized as key features of the physiological roles of these receptors.^1–5^ The ability to control the spatiotemporal profile of signaling, *i.e.*, both the subcellular location and the duration of signaling, is important for understanding of GPCR function. Sustained signaling has been proposed as an avenue towards enhanced drug efficacy.^6^ The GLP-1R, an archetypal class B1 GPCR that is a highly validated target for the treatment of type 2 diabetes and obesity,^7,8^ responds to the native agonist, GLP-1(7-36)NH2, by trafficking to membrane nanodomains,^9^ undergoing rapid internalization^10–13^ and signaling from both plasma membrane and endosomal compartments.^14^ Synthetic analogues of GLP-1 and exendin-4 with altered spatiotemporal signaling profiles have been reported.^15,16^ The importance of sustained agonist engagement with the receptor has been explored with an exendin-4 analogue that can bond covalently to a SNAP-tagged receptor and subsequently detach upon addition of a reducing agent.^17^ Photoswitchable GLP- 1R agonists and positive allosteric modulators have also been used to modulate temporal patterns of signaling.^18,19^ While these approaches are valuable, they are not well-suited for exploring spatiotemporal signaling *in vivo*.

Discovery of new agonists that display altered spatiotemporal signaling profiles at the wild-type receptor relative to GLP-1 remains an important goal. Here we report an analogue of GLP-1(7-36)NH2, designated peptide **1** below, that causes distinct signaling behavior and GLP-1R trafficking relative to the native hormone, including prolonged cAMP production in the absence of receptor internalization. Peptide **1** differs from GLP-1(7-36)NH_2_ at two positions: Val-16 is replaced by α-aminoisobutyric acid (Aib), and Ser-18 is replaced by a cyclic β-amino acid residue derived from (*S,S*)-*trans*-2- aminocyclopentanecarboxylic acid (ACPC; Figure 1A). To gain insight on the factors necessary for the unique signaling profile displayed by peptide **1**, we examined several related peptides. The analogue of **1** that contains only ACPC substitution at position 18 of GLP-1 (designated peptide **2** below) is similar to **1** in causing prolonged signaling duration at the GLP-1R, but **2** differs from **1** in causing receptor internalization comparable to that induced by GLP-1 itself. Thus, choice of residue identity at position 16 and/or 18 provides distinct and versatile spatiotemporal control over GLP-1R signaling.

**Figure 1:**
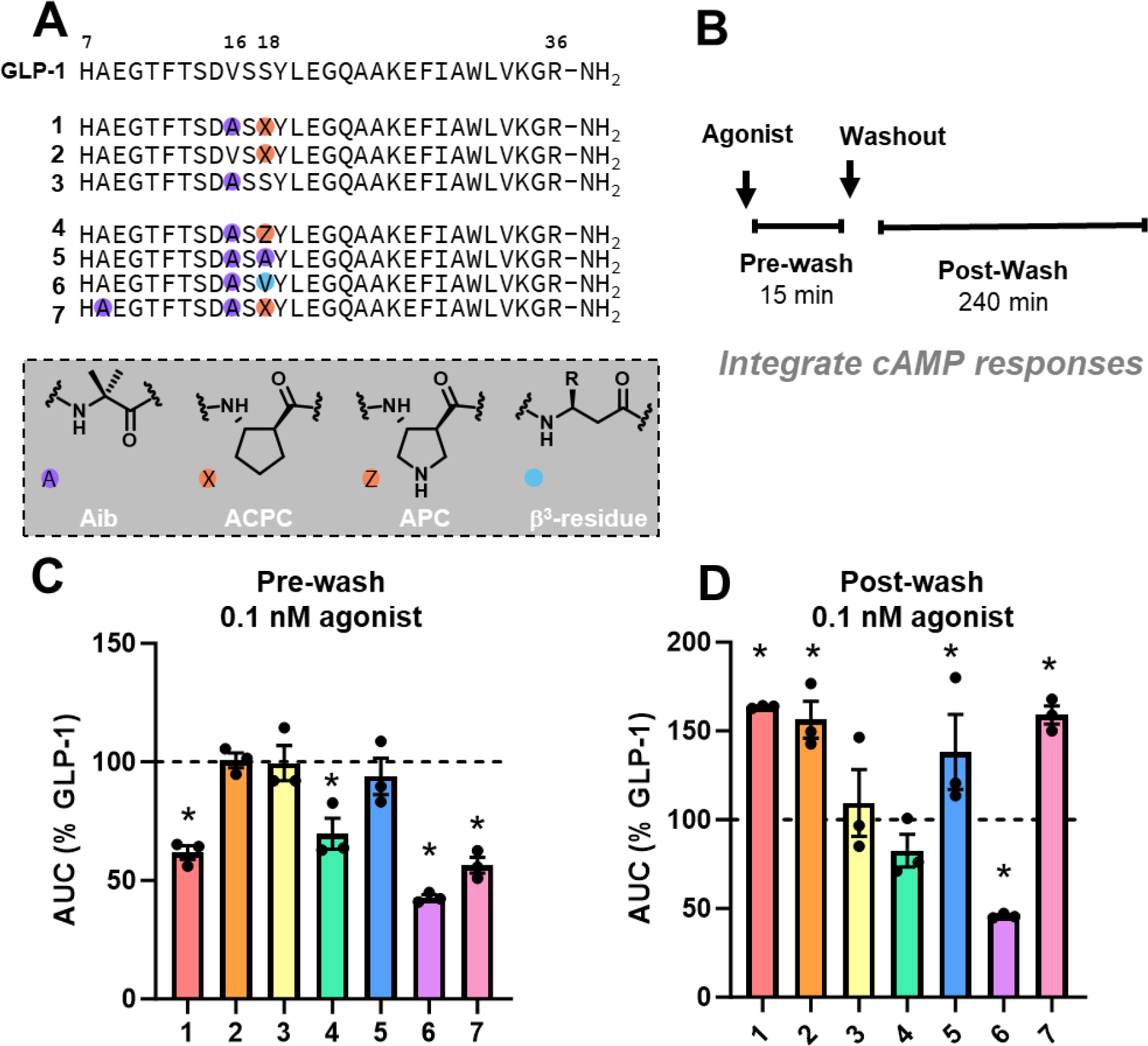
cAMP washout experiments. (**A**) Top: Sequences of GLP-1 and analogues. Select residue numbers are shown above. Bottom: Structures of non-native residues. b^3^-residues share the same sidechain (depicted as R) as the native residue indicated inside the blue circle. (**B**) Depiction of the washout assay protocol. HEK293 GS22 cells transiently expressing hGLP-1R were challenged with agonist for 15 min (pre-wash), washed, and then luminescence was monitored for 4 hours (post-wash). (**C**) AUC values for 0.1 nM agonist pre-wash (**D**) AUC values for 0.1 nM agonist post-wash. n = 3. Washout assay results where data were not normalized to GLP-1 (100%), for which statistical tests were performed (* indicates P < 0.05, one-way ANOVA with Dunnett’s post-test), are available in Figure S1 and Table S1. Error bars are standard error of the mean.

Cryo-electron microscopy (cryo-EM) structures of GLP-1R:Gs complexes containing peptides **1** and **2** offer new insight into the role of dynamics in GLP-1R activation. These data raise the possibility that the receptor engagement mode required for an agonist to stimulate cAMP production via the GLP-1R may differ significantly from the receptor engagement mode required to induce GLP-1R internalization.

## Results

### Peptide 1 induces prolonged signaling relative to GLP-1

The two sequence modifications found in GLP-1(7-36)NH_2_ derivative **1** (Figure 1A) led to a ∼5-fold decrease in potency for stimulating cAMP production in cells overexpressing the GLP-1R relative to GLP-1(7-36)NH_2_ (henceforth referred to as “GLP-1”), as quantified in Table 1. A previous study found a similar modest increase in EC_50_ when the same two modifications were implemented in a GLP-1(7-37)NH_2_ derivative, relative to GLP-1(7-37)NH_2_ (spatiotemporal features of signaling were not evaluated in the prior study).^20^

**Table 1:** EC_50_ values and maximum responses from three-parameter sigmoidal fits for concentration- response data in Figure 2A, as well as mean responses from data in figures 1C, 1D and 2D normalized to GLP-1 (100%). * indicates P < 0.05. P-values were calculated using one-way ANOVA with Dunnett’s post-test compared to GLP-1. For the washout and receptor internalization assays, statistical tests were performed on values that were not normalized to GLP-1, and these values are shown in Table S1. Uncertainties are expressed as standard error of the mean. n = 3, except for receptor internalization for peptides 2 and 3, where n = 2.

|  | cAMP Production (15 min) |  |  | Washout Assay |  | Receptor Internalization |
| --- | --- | --- | --- | --- | --- | --- |
|  | pEC<sub>50</sub> | EC<sub>50</sub> (nM) | % Max | Pre-wash AUC (% GLP-1) | Post-wash AUC (% GLP-1) | % GLP-1 |
| **GLP-1** | 10.7 ± 0.05 | 0.022 | 100 ± 2 | 100 | 100 | 100 |
| **1** | 9.96 ± 0.08* | 0.11 | 97 ± 3 | 62 ± 3 | 164 ± 1 | 2.9 ± 2 |
| **2** | 10.5 ± 0.07 | 0.031 | 101 ± 2 | 100 ± 3 | 156 ± 10 | 84 ± 2 |
| **3** | 10.5 ± 0.07 | 0.029 | 102 ± 2 | 99 ± 7 | 109 ± 19 | 96 ± 1 |
| **4** | 10.2 ± 0.06 | 0.067 | 104 ± 2 | 70 ± 9 | 83 ± 9 | 17 ± 4 |
| **5** | 10.6 ± 0.1 | 0.028 | 112 ± 4 | 94 ± 10 | 138 ± 21 | 88 ± 10 |
| **6** | 9.66 ± 0.07* | 0.22 | 102 ± 3 | 43 ± 3 | 46 ± 1 | 0.6 ± 7 |
| **7** | 9.86 ± 0.07* | 0.14 | 95 ± 3 | 56 ± 5 | 159 ± 5 | 2.8 ± 6 |

We undertook washout assays^21–24^ to examine duration of signaling by GLP-1, peptide **1** and related peptides. Cells expressing the GLP-1R were challenged with agonist, and intracellular cAMP production was monitored over time with the GloSensor luminescent biosensor system;^25^ after the maximal cAMP signal was reached (∼15 min), the agonist-containing medium was removed, the cells were washed, fresh medium was added, and the decay of the cAMP signal was monitored (Figure 1B). When 0.1 nM agonist was used (near the EC_50_), peptide **1** displayed a lower pre-wash area under the curve (AUC_15min_) response compared to GLP-1, consistent with the lower observed potency of peptide **1** relative to GLP-1 noted above. However, peptide **1** showed a significantly higher *post-wash* AUC_240min_ response relative to GLP-1 at 0.1 nM, indicating increased signaling duration for peptide **1** (Figure 1D, S1, Table S1).

We evaluated the impact of each of the two substitutions in peptide **1** individually, relative to GLP- 1, by examining peptides **2** (ACPC at position 18 only) and **3** (Aib at position 16 only). Both peptides with single substitutions were indistinguishable from GLP-1 in potency for stimulating cAMP production (Figure 2A, Table 1), and both matched GLP-1 in terms of pre-wash (AUC_15min_) response at 0.1 nM (Figure 1C, S1, Table S1). However, peptide **2** showed a prolonged post-wash (AUC_240min_) response relative to GLP-1 (Figure 1D), indicating that ACPC at position 18 is sufficient for increased signaling duration. In contrast, peptide **3**, substituted only with Aib at position 16, was indistinguishable from GLP-1 in terms of post-wash (AUC_240min_) response.

**Figure 2:**
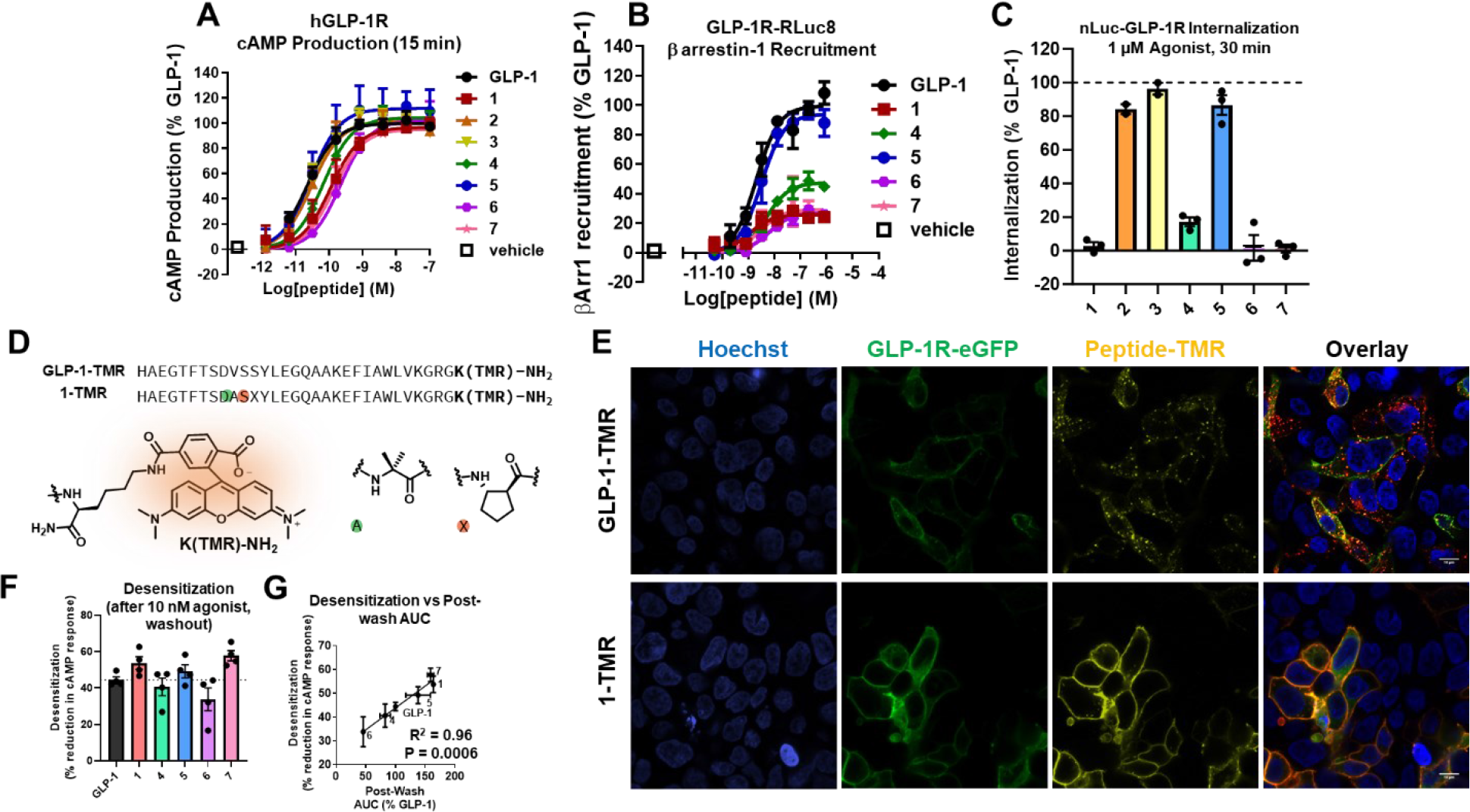
(**A**) Single time point (∼15 min post-ligand addition) maximal, concentration-response cAMP production in HEK293 GS22 cells transiently transfected with hGLP-1R. (**B**) Concentration-response curves showing recruitment of ß arrestin-1 to GLP-1-RLuc8 in transiently transfected HEK239FT cells (∼45 min post-ligand addition). (**C**) Receptor internalization as measured by reduction in maximal luminescence in HEK293 cells expressing nLuc-GLP-1R. 0% internalization was determined by cells treated with vehicle. Error bars indicate standard deviation. n = 3 except for peptides 2 and 3 which are n = 2. Internalization assay data that have not been normalized to GLP-1 (100%), for which statistical tests were performed, are available in Table S1. (**D**) Sequence of 6-carboxytetramethylrhodamine (TMR) labeled GLP-1 analogues. (**E**) Representative (n = 2) confocal microscopy images of HEK293 GS22 cells transiently expressing GLP-1R-eGFP. Cells were treated with 10 nM agonist for 30 min before washing and paraformaldehyde fixation. Top: cells treated with 10 nM GLP-1-TMR Bottom: cells treated with 10 nM 1-TMR. From left to right: Nuclear staining with Hoechst, GLP-1R-eGFP, TMR labeled peptide and overlay of the three channels. The scale bars indicate 10 µm. (**F**) (Left) Desensitization assay where HEK293 GS22 cells transiently transfected with hGLP-1R were treated with the indicated peptide for 15 min, followed by a washout step with the cells were incubated for 4 h, and then the cells were rechallenged with 100 nM GLP-1. Values indicate percent loss in cAMP signal compared to vehicle treatment. (**G**) Desensitization vs Post-wash AUC values (from Figure 2D). The line indicates a linear regression, and the P value was calculated from an F test. Error bars represent standard error of the mean unless otherwise stated.

### Amino acid residue structure at position 18 is important for the prolonged signaling of peptide 1

To probe the features of the residue at position 18 in peptide **1** that cause enhanced signaling duration, we examined peptides **4-6** (Figure 1A). Peptide **4** contains a heterocyclic analogue of the cyclic β residue ACPC (APC, structurally depicted in Figure 1A). This modification maintains the conformational constraint provided by the five-membered ring while eliminating the hydrophobicity of ACPC, because the ring nitrogen of APC should be protonated and therefore cationic near neutral pH. Both ACPC and APC promote α-helix-like secondary structure.^26,27^ To examine the role of enhanced helix propensity at position 18 in the absence of a β residue, we synthesized peptide **5**, which contains Aib at position 18, in addition to the Aib at position 16.^28^ To examine the role of helix propensity at position 18 with retention of hydrophobic α-to- β substitution, we synthesized **6**, in which the β residue lacks a cyclic constraint; acyclic β residues have diminished helical propensity relative to ACPC or APC.^26,27^ Peptide **5** had similar potency to GLP-1 in stimulating cAMP production, while peptides **4** and **6** were ∼3-fold and 10-fold less potent, respectively (Figure 2A; Table 1).

Washout assay comparisons of peptides **1** and **4-6** highlighted the importance of a helix-promoting constraint and hydrophobicity at position 18 for increased signaling duration relative to GLP-1 (Figure 1D). After stimulation with 0.1 nM agonist, cAMP production induced by peptide **5** did not differ significantly from GLP-1 pre-wash but was moderately increased post-wash relative to GLP-1 (Figure 1C, 1D, S1, Table S1). Peptide **6** displayed significantly lower AUC values for cAMP production pre- and post-wash (Figure 1C, 1D, S1, Table S1). Peptide **4** showed a moderately lower pre-wash cAMP AUC, yet a similar post-wash AUC compared to GLP-1. Because ACPC is hydrophobic, substitution of ACPC for the native Ser- 18 in peptides **1** and **2** might increase non-specific binding to surfaces or cell membranes and prevent efficient washout. This possibility seems unlikely, however, given the lack of prolonged signaling observed for peptide **6**, which has a hydrophobic but flexible β residue at position 18; reverse-phase chromatography retention times suggest that **6** is more hydrophobic than **1** (Figure S2).

We extended our evaluation to peptide **7**, which contains both substitutions found in peptide **1** along with a second Aib substitution at position 8 (Figure 1A), because of potential utility for future *in vivo* studies. The modification near the N-terminus blocks peptide cleavage by dipeptidyl peptidase 4, a process that very rapidly inactivates GLP-1 *in vivo*.^29^ In the context of other GLP-1R agonist sequences, placement of Aib at position 8 has minimal impact on activity.^29–31^ Indeed, peptide **7** was indistinguishable from **1** in terms of potency for inducing cAMP production and pre- and post-wash AUC at 0.1 nM (Figure 1D, 2A, Table 1, S1); thus, peptide **7** displays enhanced duration of signaling relative to GLP-1.

### Peptide 1 induces decreased β-arrestin-1 recruitment relative to GLP-1

β-arrestins, intracellular proteins that bind to activated GPCRs,^32^ have disparate effects on agonist-induced cAMP production, depending on the receptor. siRNA knockdown of β-arrestins-1 and -2 led to decreased cAMP production at the parathyroid hormone receptor-1 (PTHR1)^33^ but increased cAMP production at the β2-adrenergic receptor.^34^ β-arrestin-1 knockdown led to a reduction in GLP-1-mediated cAMP production in rodent insulinoma cells.^35^ Differential recruitment and activation of partner proteins such as β-arrestins might influence the duration of signaling induced by modified GPCR agonists.^21,33^ Therefore, we assayed the GLP-1 analogues with modifications at positions 16 and 18 for the ability to induce β-arrestin-1 recruitment to the GLP-1R via a BRET experiment (Figure 2B).^20^

Peptides **1** and **4-7** are potent inducers of β-arrestin-1 recruitment to the GLP-1R (similar EC_50_ values); however, most of these peptides had reduced maximal recruitment compared to GLP-1 (Table S2). The results for **1** are consistent with an earlier study,^20^ which found that the analogue of GLP-1(7-37)NH_2_ containing Aib at position 16 and ACPC at position 18 displayed a ∼70% reduction in the β-arrestin-1 recruitment maximum but no loss in potency relative to GLP-1(7-37)NH_2_ (Figure 2B, Table S2). Peptide **5**, which contains Aib at position 18 and was the most potent analogue in the cAMP assay, was comparable to GLP-1 in β-arrestin-1 recruitment maximum, but the analogues containing a β residue at position 18 (peptides **1**, **4**, **6** and **7**) had β-arrestin-1 recruitment maxima that were only 25-50% that of GLP-1. We note that peptide **6** was comparable to **1** in terms of a low β-arrestin-1 recruitment maximum (Table S2) and in potency for stimulating cAMP production (Table 1), but that there was a sharp contrast between **1** and **6** in terms of duration of signaling (Figure 1D). This comparison between peptides **1** and **6** suggests that lower β-arrestin-1 recruitment is not sufficient for enhanced duration of cAMP signaling.

Analysis with the Black-Leff operational model of agonism^36,37^ indicated that none among peptides **1** and **4-7** was significantly biased towards either cAMP production or β-arrestin-1 recruitment relative to GLP-1 (Table S2). While agonists that are biased away from β-arrestin recruitment can exhibit increased signaling duration,^21,38^ the behavior of peptide **1** suggests that this type of bias is not required for prolonged stimulation of cAMP production via the GLP-1R.

### Peptide 1 induces prolonged signaling at the cell surface

Activation of the GLP-1R by the native hormone promotes G protein-mediated signaling at the cell surface and in endosomes.^14,39^ We examined whether the prolonged signaling observed for peptide **1** is associated with a particular subcellular location by using a version of the GLP-1R that has nanoluciferase (nLuc) fused to the N-terminus^40^ to monitor internalization in response to stimulation by an agonist. nLuc-GLP-1R internalization is implicated by a decrease in luminescence due to the reduced availability of luciferase substrate inside the cell versus at the cell surface.^41^ Luciferase substrate was introduced 30 min after addition of excess (1 µM) GLP-1 or peptides **1-7**. As expected, after GLP-1 treatment a substantial decline in luminescence was observed, which is consistent with extensive nLuc-GLP-1R internalization induced by the native hormone (Figures 2C, S3A). In contrast, no internalization was detected after treatment with peptide **1**, **6** or **7** (Figure 2C, Table 1, S1). Peptide **4** exhibited substantially reduced receptor internalization relative to GLP-1. Only peptide **5** among the doubly substituted analogues was comparable to GLP-1 in terms of receptor internalization.

The lack of receptor-internalization activity observed for peptide **1** provides a striking contrast to the behavior of singly substituted peptides **2** and **3**, which both were comparable to GLP-1 in their ability to induce GLP-1R internalization (Figure 2C, Table 1, S1). Thus, the ACPC-18 and Aib-16 substitutions act in concert to prevent receptor internalization upon engagement of peptide **1**. These data suggest that peptide **1** induces prolonged signaling from receptors that remain at the cell surface and that peptide **7** behaves similarly. Although peptide **2** induces prolonged signaling, in this case the receptor is internalized.

To test the conclusion that peptide **1** induces signaling exclusively from receptors at the cell surface, we evaluated localization of this peptide via microscopy. These experiments employed derivatives of GLP- 1 and peptide **1** labeled with the fluorescent dye, 6-carboxytetramethylrhodamine (TMR), linked by an amide bond to a lysine side chain at the peptide C-terminus (Figure 2D). In preliminary studies, we evaluated the abilities of **GLP-1-TMR** and **1-TMR** to activate the GLP-1R. The TMR-labeled analogues showed nearly indistinguishable EC_50_ values for cAMP production relative to the unlabeled counterparts (Figure S3B, Table S3). Thus, placing the TMR fluorophore at the C-termini of these peptides does not alter agonist function, as monitored by cAMP production. For the microscopy studies, we used a version of the GLP-1R that has eGFP fused to its C-terminus (GLP-1R-eGFP).^42^

HEK293 cells transiently expressing GLP-1R-eGFP were treated with either **GLP-1-TMR** or **1- TMR** (10 nM) for 30 min, washed, fixed with paraformaldehyde, and imaged with confocal microscopy. **GLP-1-TMR** showed predominantly punctate distribution, which is consistent with location of this peptide in endosomes. In contrast, **1-TMR** largely colocalized with GLP-1R-eGFP at the cell surface (Figure 2E). These observations support the conclusion that peptide **1** binds to the GLP-1R but fails to induce receptor internalization.

### Peptide 1 is comparable to GLP-1 in decreasing cellular response to GLP-1 restimulation

β- arrestin-1 recruitment and receptor internalization are classically associated with GPCR signal termination and a decline in cellular response to subsequent stimulation.^43^ Thus, our agonists with reduced β-arrestin-1 recruitment maxima and reduced induction of receptor internalization relative to GLP-1 (peptides **1**, **4**, **6** and **7**) could be less effective than GLP-1 at causing a diminished cellular response to restimulation. To examine this hypothesis, we performed washout assays with 10 nM agonist, and after 4 h, we restimulated the cells with a saturating concentration of GLP-1 (100 nM; Figure 2F, S4, Table S2). Our hypothesis predicted that cells initially stimulated by peptides **1**, **4**, **6** or **7** would show a greater response to the second phase of stimulation relative to cells initially stimulated by GLP-1; the data described below invalidated this hypothesis.

In a positive control study with cells initially stimulated with GLP-1, we observed restimulation with GLP-1 caused only ∼60% cAMP production relative to initial treatment with vehicle. This observation is consistent with the expectation that the initial GLP-1 treatment induced loss of cAMP responsiveness.

Pretreatment with peptide **1** led to a significant decrease in cAMP production following subsequent stimulation with GLP-1, relative to pretreatment of the cells with vehicle (Figure S4, Table S2); the extent of the decrease was similar to that observed for pretreatment with GLP-1. This observation, and the microscopy data indicating that the GLP-1R remains at the cell surface after engagement by peptide **1** (Figure 2E), suggest that the decline in cAMP production upon restimulation with GLP-1 in this case involves a mechanism other than receptor internalization. Previous studies have suggested that phosphorylation of the C-terminal tail of the GLP-1R can cause loss of cAMP responsiveness that is independent of receptor internalization.^44,45^ This mode of desensitization was previously attributed to recruitment of β-arrestin,^46^ but our observations raise the possibility of a different mechanism, since maximal β-arrestin recruitment is low for peptide **1**.

Peptides **4-7** were similar to GLP-1 and peptide **1** in that following an initial cellular treatment with peptides **4-7**, cAMP production was decreased during subsequent stimulation with GLP-1, relative to initial treatment of the cells with vehicle. Among the peptides we examined, the decreases in cAMP responsiveness were well-correlated (R^2^ = 0.96, P = 0.0006) with integrated cAMP responses from washout experiments (Figure 2G). This correlation suggests that cAMP itself might be a key determinant of the extent to which cells respond to a second agonist stimulation, which represents a contrast with the prior observation that cAMP does not play a role in GLP-1R desensitization.^45^ Desensitization mediated by cAMP is precedented for a different GPCR: the β2-adrenergic receptor is phosphorylated by protein kinase A, an enzyme that is activated by cAMP, and receptor phosphorylation diminishes response to agonist.^47^ Our data raise the possibility that a similar mechanism may be at work for the GLP-1R; further studies will be required to test this hypothesis.

### Cryo-EM structures of 1 and 2 bound to the GLP-1R

To elucidate the interactions of **1** and **2** with the GLP-1R, we performed single-particle cryo-EM studies of complexes in which each peptide was bound to the receptor engaged by a G protein heterotrimer. We expressed the GLP-1R, dominant negative Gαs,^48,49^ Gβ1, and Gγ2 in insect cells. We then added nanobody35 and saturating peptide to form a complex, solubilized membrane proteins in lauryl maltose neopentyl glycol with cholesteryl hemisuccinate, and isolated the complex via anti-FLAG affinity.^40,50,51^ Further purification via size-exclusion chromatography provided homogenous samples (Figure S5).

After vitrification, the complexes were imaged with a Glacios 200 kV transmission electron microscope.^52^ From these datasets, consensus maps of the complexes containing peptide **1** or **2** bound to GLP-1R/Gs were refined to 3.44 Å and 2.97 Å (Figures 3A, S6-S8, Table S4), respectively, enabling modeling for most components in each complex. In both cases, the Gs α-helical domains and sections of ICL3 were poorly resolved, which precluded modelling of these portions. The receptor core and G protein were similar in the complexes containing peptides **1** or **2**, and these features were similar to their counterparts in the complex of GLP-1 bound to GLP-1R/Gs.^53^ In contrast, the peptide-binding pockets and extracellular domains for the complexes containing peptides **1** or **2** bound differed from the analogous portions of the complex containing GLP-1.

**Figure 3:**
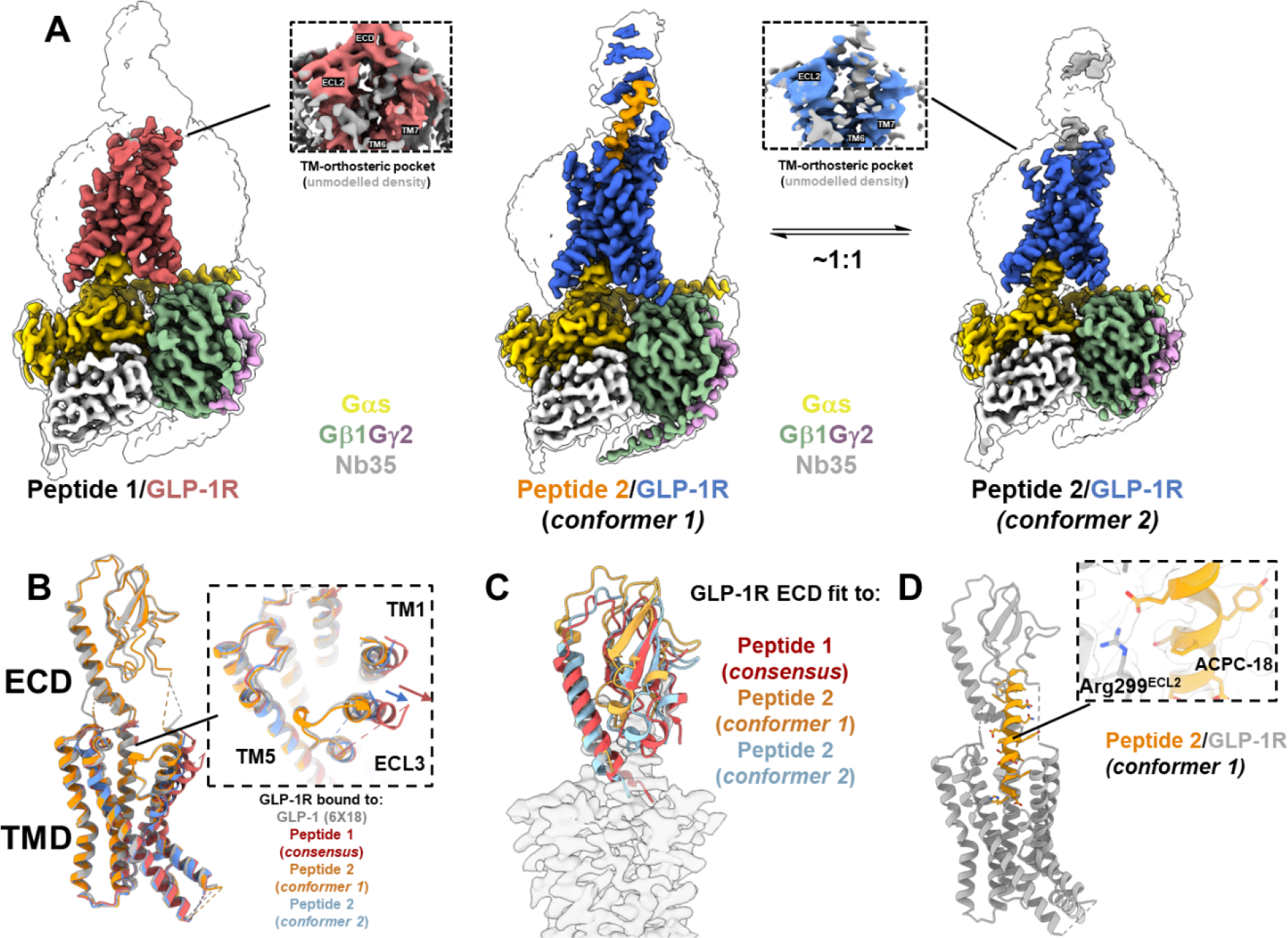
(**A**) EM density maps of peptides **1** and **2** bound to GLP-1R/Gs complexes. The transparent map with a black silhouette is contoured at low threshold and colored maps are contoured at higher threshold. (**B**) A comparison of the models of the GLP-1R bound to GLP-1 (PDB: 6X18), peptide 1, peptide 2 (conformer 1) and peptide 2 (conformer 2) shown in gray, red, orange and blue, respectively. Peptide ligands were removed for clarity. The extracellular domains for peptide 1 and peptide 2 (conformer 2) were not modelled due to low local resolution. (**C**) The crystal structure of the GLP-1R extracellular domain (PDB: 3IOL) rigidly fit (using Chimera 1.14) to each receptor-focused cryo-EM density maps indicated by each color. The aligned EM map of peptide 2 conformer 1 is shown in silhouette for reference. (D) The structure of peptide 2 bound to GLP-1R in conformer 1.

### Peptide 1 is highly dynamic and induces a distinctive extracellular domain conformation when bound to the GLP-1R

While the global resolution for the cryo-EM map of the complex containing peptide **1** was resolved to 3.44 Å, local resolutions for the peptide and the extracellular domain (ECD) were substantially lower (Figure S8), indicating high flexibility in these components relative to the rest of the complex. 3D-variability analysis^54^ and 3DFlex refinement^55^ partially resolved the conformational heterogeneity (Figure S8, Video S1), but these methods were unable to completely recover high-resolution spatial information. Poor local resolution prevented us from confidently modeling the peptide at the atomic level or directly visualizing the interactions of Aib-16 or ACPC-18 with the GLP-1R. Despite the poor resolution of bound peptide **1**, there was clear density indicating peptide engagement with the ECD (Figure S9). The cryo-EM data allowed us to detect outward positioning of extracellular-loop 3 (ECL3) and transmembrane-helix 7 (TM7) relative to the complex containing GLP-1 and the GLP-1R (Figure 3B). Outward positioning of ECL3 and TM7 similar to those observed in our structure with **1** has been observed with several other agonists,^40,56^ notably with cAMP-biased agonists, but the mechanistic implications of these conformational changes are not obvious given the disparate activity profiles of these agonists.

Local refinement of the receptor density provided a map of relatively low resolution (Figure S8), but the map was sufficient to enable rigid docking of the ECD crystal structure into density corresponding to the ECD (Figure S9).^57^ The binding of peptide **1** induces a distinct consensus orientation of the ECD relative to the transmembrane domain (TMD) compared to the orientation observed when GLP-1 is bound (Figure 3C). In the complex containing **1**, there is a ∼14° rotation of the ECD relative to the TMD and an ∼8 Å movement of the ECD N-terminal α-helix towards the TMD (Figure 3C, S10). Because the ECD plays a critical role in most peptide-mediated activation of the GLP-1R,^58^ the distinctive ECD-TMD juxtaposition observed in the complex containing peptide **1** might influence receptor activation. We hypothesize that the native hormone can induce receptor conformations similar to those captured in the complex with peptide **1** during the activation process, but that for GLP-1 these states are too transient to be detected via cryo-EM. Additionally, we hypothesize that a state in which peptide **1** adopts a position similar to GLP-1 is possible, but the substitutions at positions 16 and 18 and the absence of stabilizing interactions of peptide **1** with TM6/ECL3 make this mode of peptide-receptor engagement sufficiently unfavorable and transient to preclude visualization via cryo-EM.

### Peptide 2 exhibits two conformers when bound to GLP-1R

Heterogenous refinement of the 2.97 Å consensus map for GLP-1R/Gs bound to peptide **2** delineated two conformers that each represented approximately half of the particle projections (Figure 3A, S7). These two conformers (henceforth referred to as conformer 1 and conformer 2) were refined independently to global resolutions of 3.10 Å and 3.26 Å, respectively. The conformational heterogeneity in the GLP-1R/Gs+peptide **2** complex is reminiscent of heterogeneity in the structure of GLP-1R/Gs engaged by an exendin-4 derivative containing D-Ala in place of Gly4 (Ex4-D-Ala). This D-amino acid substitution was introduced to reduce helicity near the peptide N-terminus.^40^ Ex4-D-Ala is a potent agonist for the GLP-1R, and cryo-EM analysis of the GLP-1R complex formed with Ex-4-D-Ala revealed two distinct conformers.^40^

The structure of conformer 1 in the complex formed by GLP-1R/Gs and peptide **2** is very similar to the structure of the complex formed by GLP-1R/Gs and GLP-1.^53^ In conformer 1, the ACPC residue of peptide **2** is readily visualized in the EM density map and participates in an α-helix-like conformation (Figure 3D). Numerous crystal structures of other ACPC-containing peptides show this cyclic β residue within α-helix-like conformations.^26,59^ In the GLP-1-bound structure, Ser18 is solvent-exposed and does not form hydrogen bonds with receptor residues, so no such interactions are lost upon ACPC substitution. However, the ACPC residue in conformer 1 is close to the side chain of Arg299^ECL2^, a key receptor residue for signal transduction via cAMP.^60^ Gain or loss of non-bonded interactions with Arg299^ECL2^ upon ACPC substitution at position 18 might influence the signaling behavior that results from peptide binding, potentially including duration of signaling, relative to signaling induced by GLP-1.

The structure of conformer 2 in the complex formed by GLP-1R/Gs and peptide **2** features low resolution of the peptide, as was also observed in one of the structures that emerged from the cryo-EM analysis of Ex4-D-Ala bound to the GLP-1R.^40^ As in the prior case, the crystal structure of GLP-1 bound to the GLP-1R ECD^57^ fits the density for the peptide **2**+ECD portion of conformer 2. Helical density for peptide **2** extends from the C-terminus and appears to terminate around Thr11, which is similar to the cryo-EM data for conformer 2 in the GLP-1R+Ex-4-D-Ala complex structure (Figure S11).^40^ As with the structure of the complex containing peptide **1**, conformer 2 of the complex containing peptide **2** exhibits outward positioning and disorder of ECL3, as well as alterations to the ECD conformation (∼6° rotation in α-helix-1), relative to the complex containing GLP-1 (Figure 3C, S10).

## Discussion

We previously provided evidence that flexibility and dynamics of engagement of the N-terminal region of a peptide agonist are important for the activation of the GLP-1R^40,61–63^ despite the fact that such conformational dynamics are not directly observed in structures of the native hormone, GLP-1, bound to the GLP-1R.^53,64^ The present work strengthens the conclusion that agonist dynamics plays a significant role in GLP-1R activation, as both **1** and **2** are potent agonists, and both peptides display mobility in cryo-EM reconstructions of agonist-receptor complexes. Unlike other peptides that exhibited N-terminal mobility in previous EM data or molecular dynamics simulations,^40,61,62^ peptides **1** and **2** are identical to GLP-1 over the first 9 and 11 residues, respectively. These observations demonstrate that the mode of engagement of the agonist N-terminus with the TMD, which is presumably critical for receptor activation, can be strongly influenced by residues significantly removed from the N-terminus. Our data suggest that a single substitution toward the middle of the peptide might influence the mode and stability of binding of the peptide N-terminus to the TMD core. This explanation offers a framework for understanding previous observations that α-to-β substitutions near the center of GLP-1 can induce biased agonism,^20,31,65^ even though modification in this central region might have been expected to influence receptor affinity rather than the signaling profile.

Most reported structures of GLP-1R complexes containing peptide agonists,^40,56,61,62^ including conformer 1 of the complex containing peptide **2** that is reported here, reveal the agonist in a resolved, fully α-helical or mostly α-helical conformation. In contrast, the cryo-EM data did not reveal a similar, well-resolved α-helical conformation for peptide **1** bound to GLP-1R. Instead, the data suggest that peptide **1** binds in a highly mobile state. These observations are noteworthy given that peptide **1** contains only two substitutions relative to GLP-1, and both of the noncanonical residues introduced (Aib and ACPC) support helical secondary structure.^26,61^ The structure determined for conformer 1 of the complex containing **2** shows that ACPC at position 18 can be accommodated within a fully helical bound peptide conformation that is stably positioned relative to the receptor. Thus, the disorder of the peptide component in the complex containing peptide **1** raises the possibility that Aib at position 16 (unique to peptide **1**) forms distinctive contacts with the receptor or induces steric clashes that destabilize a complex in which the peptide adopts a well-defined helical conformation that has a well-defined position relative to the surrounding receptor surface. ACPC at position 18, alone, may cause a less profound destabilization of such a complex, given that the cryo-EM data for the complex formed by GLP-1R/Gs and peptide **2** reveal two states, one with a fully helical peptide molecule that has a well-defined binding mode and the other with a peptide that is disordered relative to the receptor and not fully helical.

Both peptides **1** and **2** contain the Ser-18 → ACPC substitution, and these two analogues showed similarly increased signaling duration compared to GLP-1 in washout assays, but a common structural feature that might explain this functional similarity was not readily apparent in the conformations derived from our cryo-EM data. It is likely that the position 18 substitution increases the conformational dynamics of the peptide when bound to the GLP-1R, as complexes containing either peptide **1** or peptide **2** exhibited a poorly resolved receptor ECD relative to the complex containing the native peptide. However, for the complex containing peptide **2**, approximately half of the particles had the peptide stably positioned relative to the receptor, while no particles of this type could be detected for the complex containing peptide **1**. The structural differences between complexes containing peptides **1** or **2** might be correlated with the functional differences between peptides **1** and **2** in terms of receptor internalization. The lack of a stably positioned agonist in the structure of the GLP-1R complex containing peptide **1** raises the possibility that a relatively static and fully helical peptide component, which has been commonly observed in structures containing other hormones bound to class B1 GPCRs,^66,67^ is important for inducing internalization of the activated receptor. However, the high agonist potency of peptide **1** suggests that receptor activation can be induced by and may even be facilitated by dynamic engagement of the agonist with the receptor.

## Conclusions

The GLP-1R, an important target for the treatment of type 2 diabetes and obesity, undergoes rapid internalization upon complexation with GLP-1, with continued cAMP production stimulated by receptors in endosomes.^11,14,39^ Here, we have established that the GLP-1 analogue peptide **1**, which contains two modifications toward the middle of the sequence, can engender prolonged stimulation of cAMP production relative to GLP-1. In addition, receptors activated by peptide **1** remain on the cell surface, which contrasts with the rapid internalization of receptors activated by the native hormone. Replacement of the native residue Ser-18 with a cyclic β-amino acid residue, ACPC, is sufficient for prolonged signaling, as shown by the behavior of mono-substituted GLP-1 analogue peptide **2**. Both substitutions in peptide **1**, Ser18→ACPC and Val16→Aib, are necessary to prevent receptor internalization. Our demonstration that the distinctive spatiotemporal receptor-activation profile of peptide **1** is preserved after an additional modification, replacement of native residue Ala-8 by Aib (peptide **7**), which blocks degradation by DPP-IV,^29^ provides a novel tool for future studies *in vivo.* Structural data for the complexes formed between peptides **1** or **2** and the GLP-1R are consistent with recent findings that dynamic engagement of the peptide agonist in the bound state is important for receptor activation.^40,62^

Our study complements and significantly extends recent reports^15,16,68–73^ in identifying GLP-1R agonist modifications that modify receptor trafficking and signaling duration. While previous studies primarily focused on cAMP production vs β-arrestin recruitment bias as a driver of altered spatiotemporal signaling profiles,^15,16,71,72^ we describe two ligands that do not appear to manifest such bias relative to GLP- 1^20^ but nevertheless display distinctive patterns of signaling and regulation. The previous studies mostly involved modifications of GLP-1 near the agonist N-terminus, whereas peptides **1**, **2** and **7** contain substitutions toward the middle of the sequence. Our findings therefore suggest that a broader exploration of analogues of GLP-1, exendin-4 and related peptides might be productive for discovery of agonists that display unique spatiotemporal signaling profiles. Agonists with novel activation characteristics, including those established here for peptides **1**, **2** and **7**, could be useful for elucidating mechanisms of signal transduction via the GLP-1R and might ultimately provide a basis for development of new therapeutic agents with tailored pharmacological properties.

## Acknowledgements

This work was supported in part by the National Institutes of Health (R01 GM056414, to S.H.G.). P.M.S. is a Senior Principal Research Fellow (#1154434) and D.W. a Senior Research Fellow (#1155302) of the National Health and Medical Research Council (NHMRC). P.M.S is the Director and D.W. the Monash University Node leader of the Australian Research Council Industrial Transformation Training Centre for Cryo-electron Microscopy of Membrane Proteins (#IC200100052). This work was funded in part by an NHMRC Program Grant (#1150083) to P.M.S and NHMRC Ideas Grant (#1184726) to D.W. B.P.C. and R.K.M. were supported in part by graduate fellowships from the US National Science Foundation (DGE-1747503). B.P.C. was supported in part by a Biotechnology Training Grant from National Institute of General Medical Sciences (NIGMS) (T32 GM008349). M.V.H. was supported in part by a Chemical Biology Interface Training Grant from NIGMS (T32 GM008505). We thank Elle Grevstad for helpful discussions and assistance with optical microscopy. Confocal microscopy was performed at the University of Wisconsin-Madison Biochemistry Optical Core, which was established with support from the University of Wisconsin-Madison Department of Biochemistry Endowment. Cryo-EM samples were imaged at the Bio21 Advanced Microscopy Facility (The University of Melbourne). High-performance computing was supported by Monash MASSIVE.

## Author contributions

B.P.C performed optical microscopy, expressed and purified the protein complexes, performed cryo-EM data processing, and built the structural models. B.P.C. and M.V.H synthesized peptides. B.P.C, M.V.H., and R.K.M. performed *in vitro* assays. M.J.B. prepared cryo-EM samples and acquired cryo-EM data. D.W., P.M.S., and S.H.G. supervised the project. B.P.C., D.W., P.M.S., and S.H.G. interpreted data and wrote the paper. All authors edited the manuscript.

## Data availability

Cryo-EM maps have been deposited in Electron Microscopy Data Bank (EMDB) under ascension codes EMD-XXXXX, EMD-XXXXX and EMD-XXXXX for GLP-1R/Gs/Peptide 1, GLP- 1R/Gs/Peptide 2 (conformer 1), and GLP-1R/Gs/Peptide 2 (conformer 2), respectively. Cryo-EM models have been deposited in the Protein Databank (PDB) under PDB IDs XXXX, XXXX, and XXXX for GLP- 1R/Gs/Peptide 1, GLP-1R/Gs/Peptide 2 (conformer 1), and GLP-1R/Gs/Peptide 2 (conformer 2), respectively.

---

**Figure S1:**
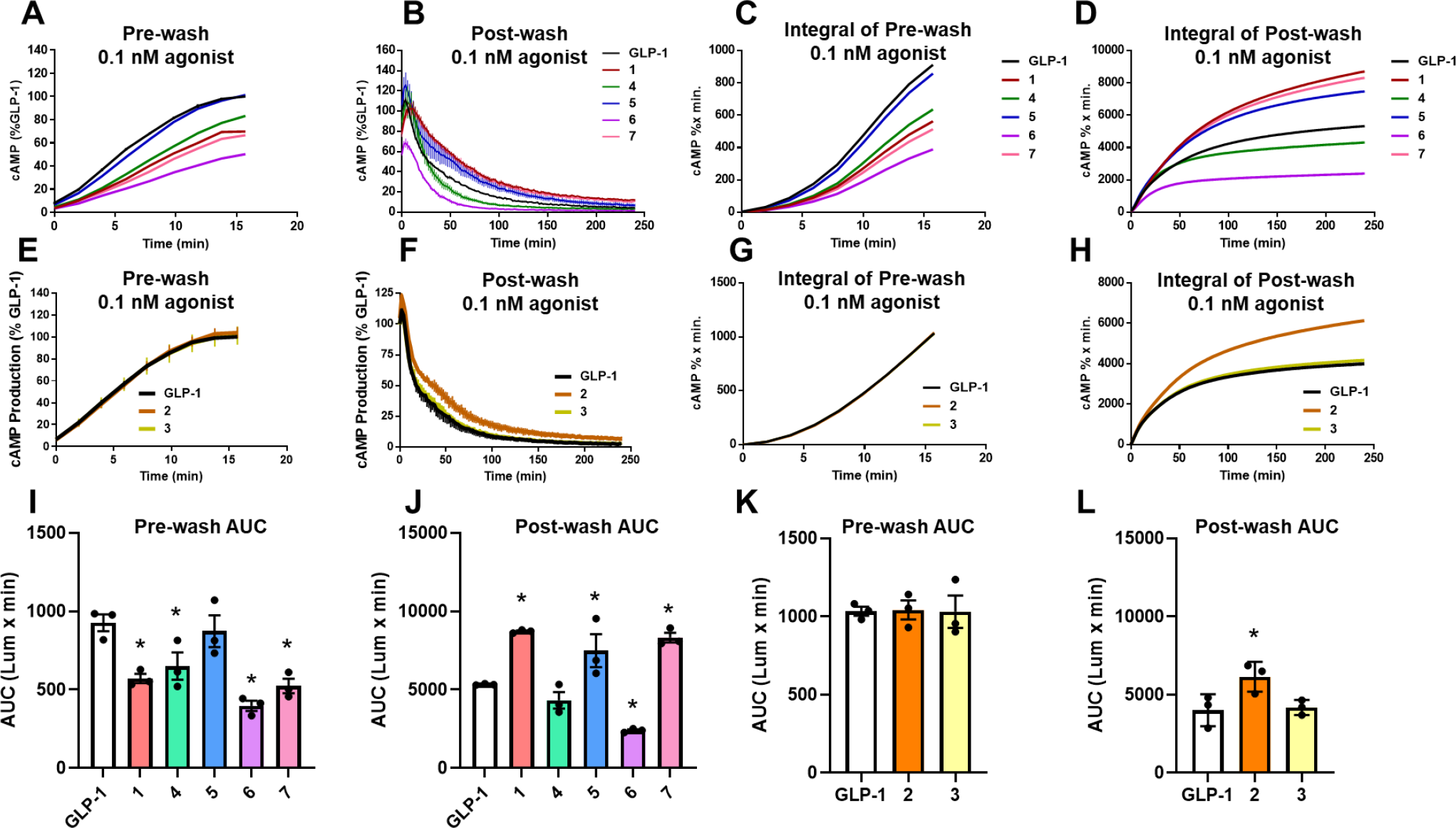
cAMP washout experiments. HEK293 GS22 cells transiently expressing hGLP-1R were challenged with agonist for 15 min, washed, and then luminescence was monitored for 4 hours. Experiments were performed in two batches of assays (each batch included GLP-1 as a control and either peptides 1,4- 7 (**A**-**D**, **I**,**J**) or peptides 2 and 3 (**E**-**H**, **K**,**L**)), which were analyzed independently by one-way ANOVA with Dunnett’s post-test compared to GLP-1. (**A,E**) 0.1 nM agonist challenge pre-wash time course (**B,F**) 0.1 nM agonist post-wash time course. (**C,G**) Integral of mean cAMP response of 0.1 nM agonist challenge pre-wash time course starting from time 0. (**D,H**) Integral of mean cAMP response of 0.1 nM agonist challenge post-wash time course starting from time 0. 100% cAMP production represents mean luminescence of GLP-1 treated cells at the final or initial timepoints for pre-wash or post-wash conditions, respectively. Error bars represent standard error of the mean. n = 3. AUC values shown in this figure were normalized to GLP-1, and these normalized values are displayed in figure 1C,1D and Table 1.

**Figure S2:**
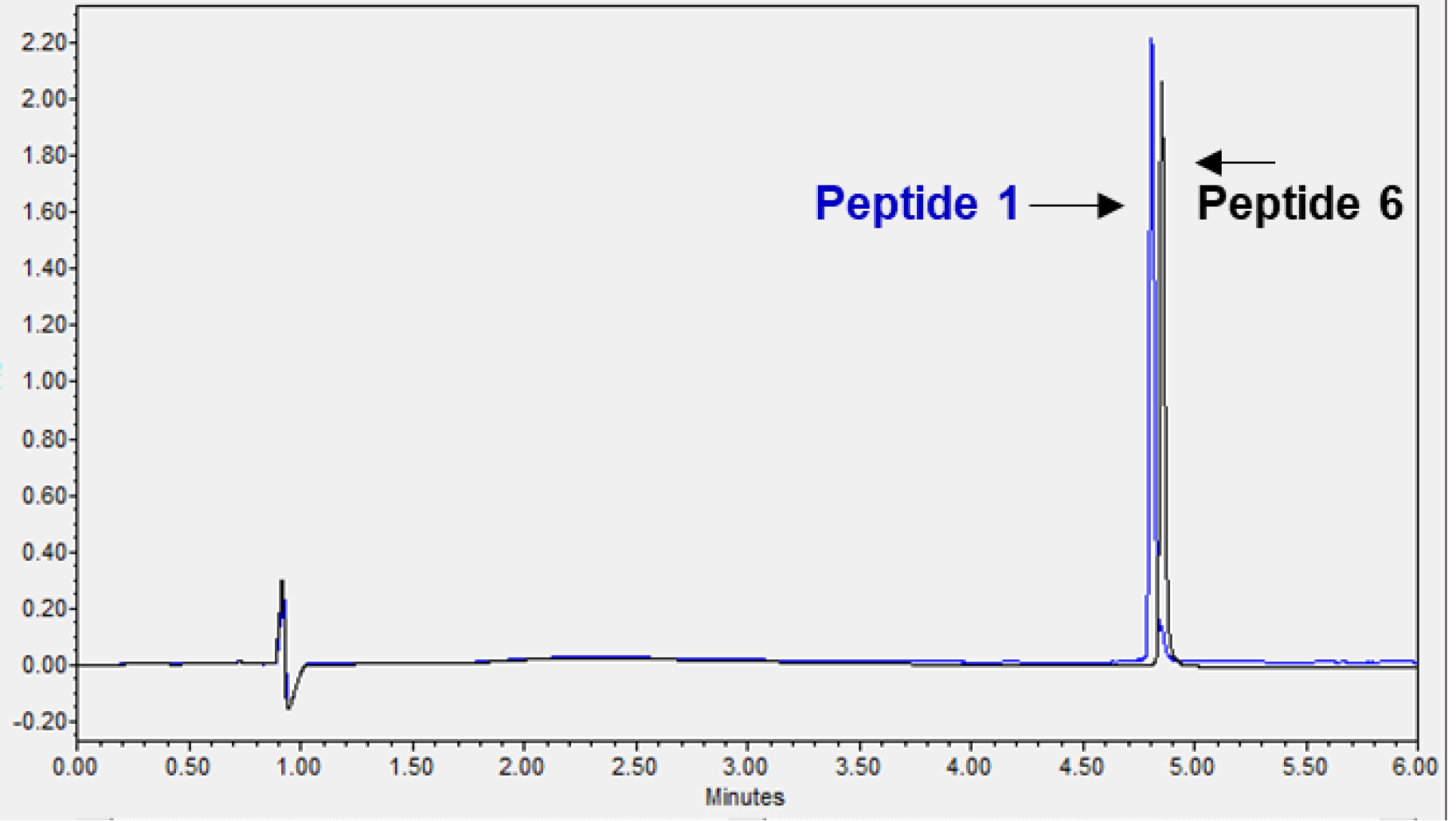
Comparison of the reverse-phased chromatography retention times of peptide 1 and 6.

**Figure S3:**
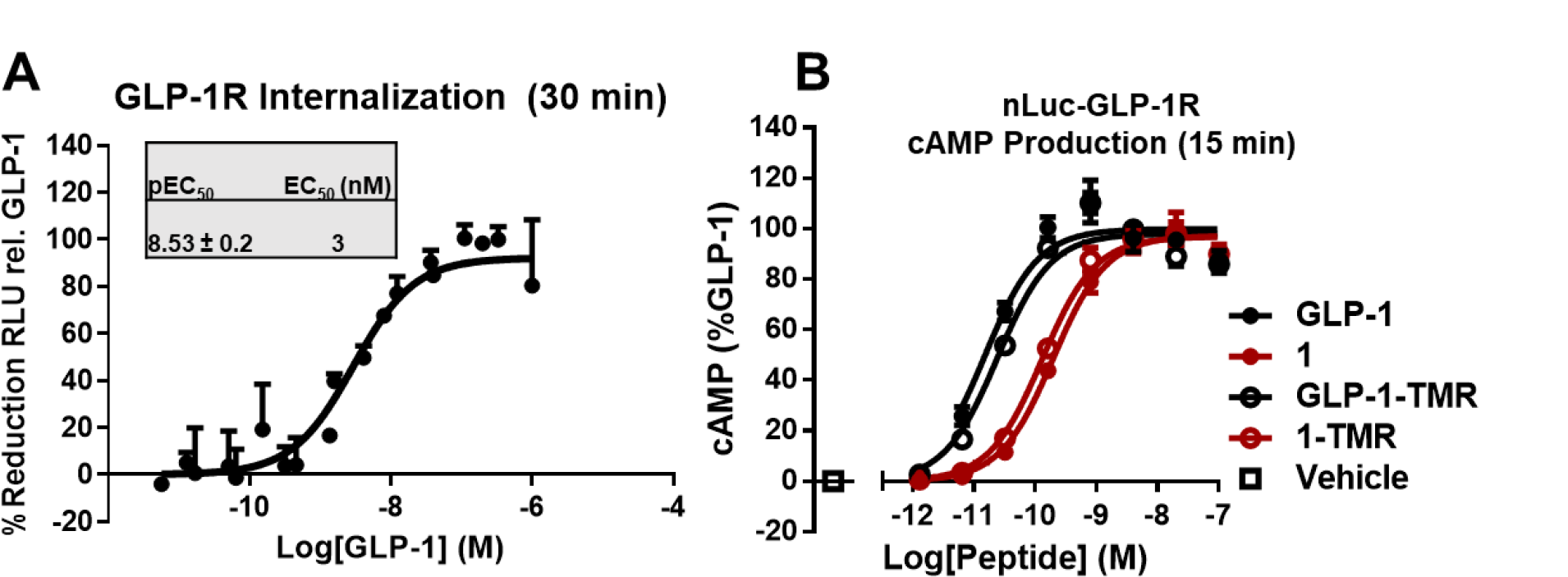
(A) Concentration-response for GLP-1 mediated receptor internalization, measured by reduction in maximal luminescence in HEK293 GS22 cells expressing nLuc-GLP-1R 30 min after ligand addition. (B) Concentration-response for cAMP production at ∼15 min post ligand addition in HEK293 cells transiently expressing nLuc-GLP-1R. Errors indicate standard error of the mean. n = 3.

**Figure S4:**
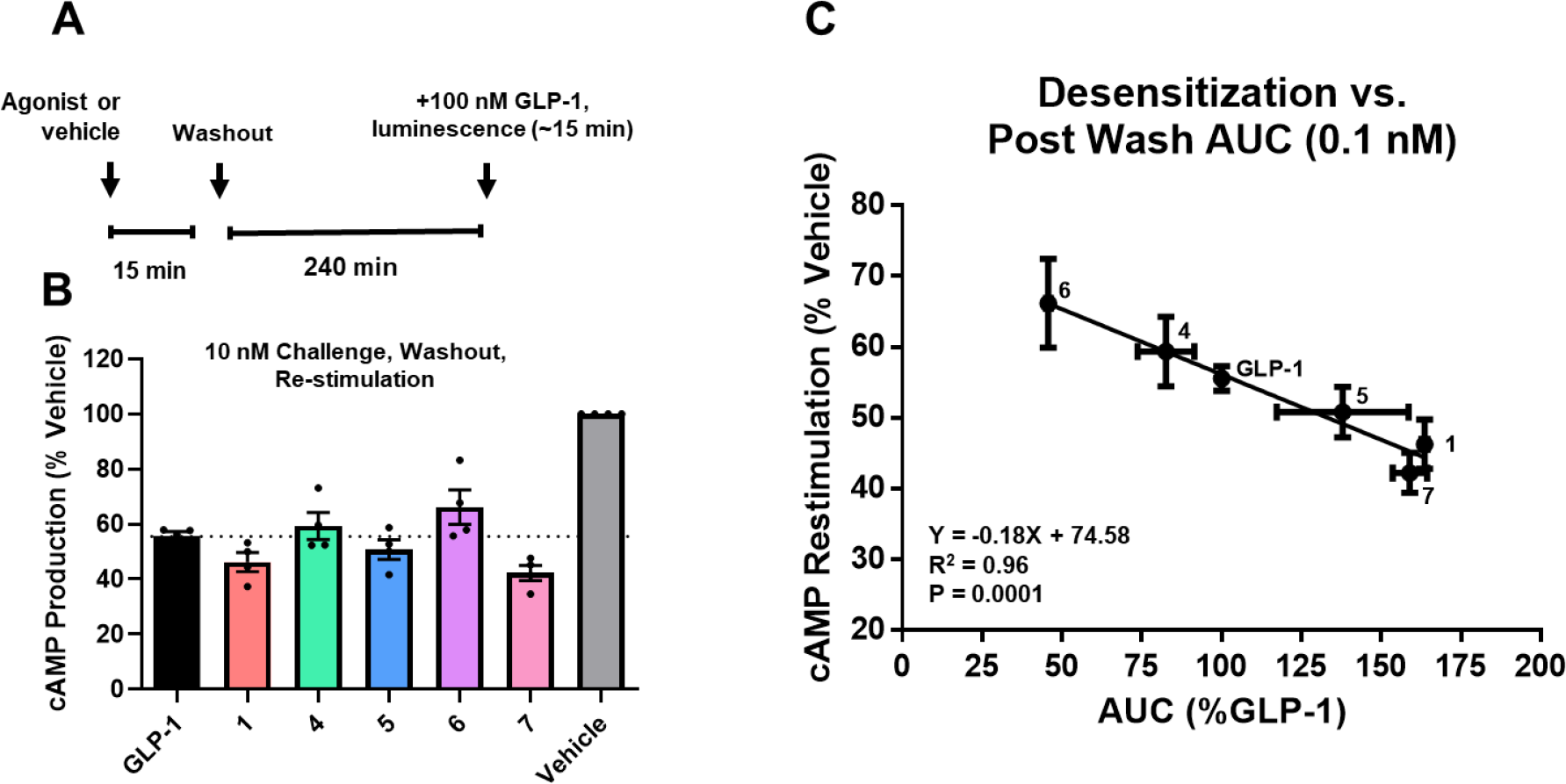
(**A**) Depiction of the desensitization assay protocol. (**B**) cAMP production in response to 100 nM GLP-1 measured 4 h after 15 min pretreatment of GLP-1R expressing HEK293 cells by vehicle (control), 10 nM GLP-1 or peptide analogues. The dotted line indicates the mean cAMP production of cells pre-treated with GLP-1. Data are normalised to the response measured in vehicle-treated cells. Lower values indicate more desensitization. (**C**) Relationship between cAMP restimulation and post-wash AUC. Datapoints are labeled with their corresponding peptide. The parameters for the best fit line, R^2^ value, and P value (F-test) are shown. N = 4 independent experiments. Error bars represent standard error of the mean. Panels **B** and **C** show data from the same experiments as shown in Figure 2F and 2G, but the data are depicted in terms of cAMP production rather than by calculated desensitization.

**Figure S5:**
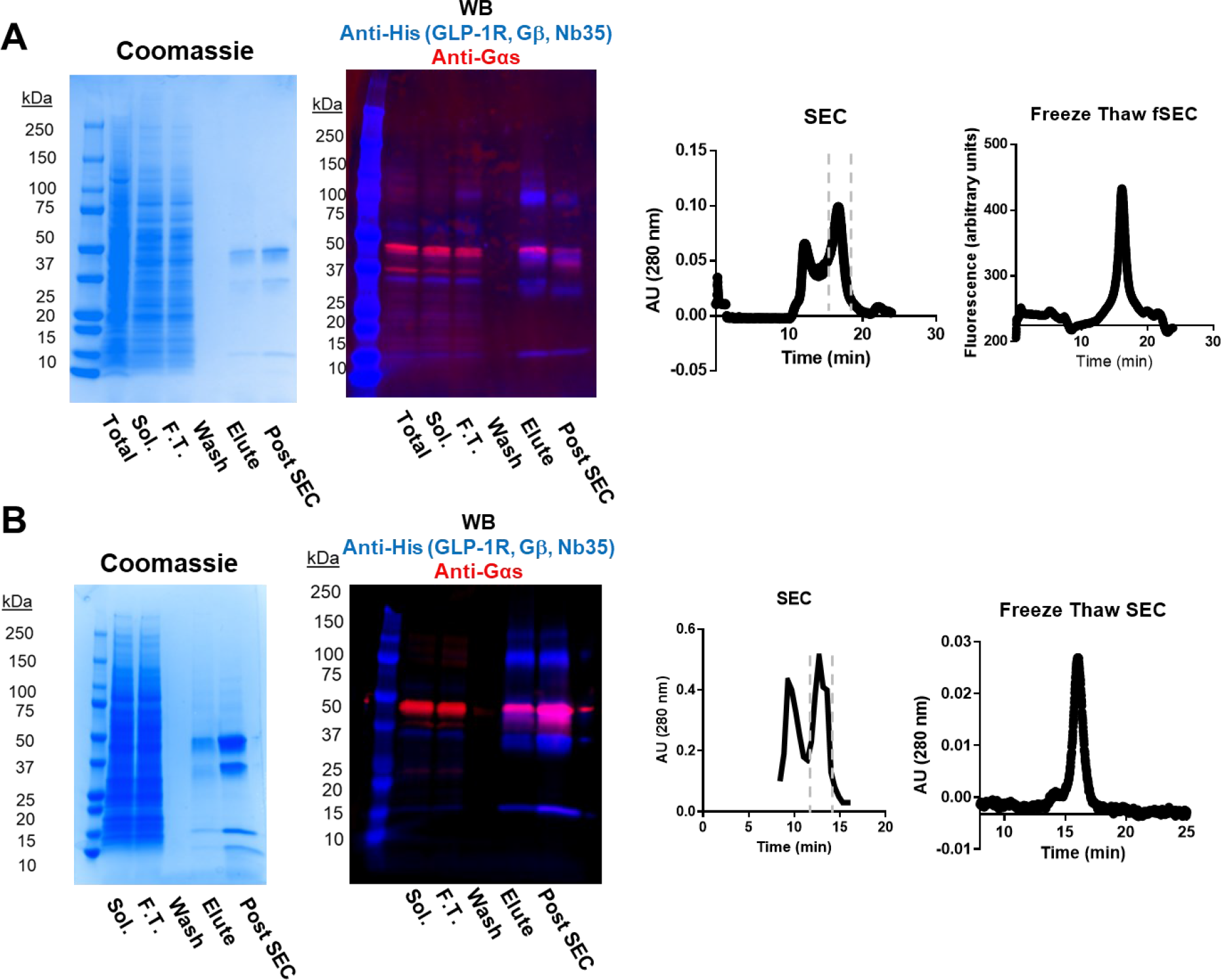
Production and characterization of GLP-1R/Gs complexes for peptides **1** (A) and **2** (B). Left: SDS-PAGE with Coomassie staining of samples. Middle-left: Western blot of complex samples. Blue coloring indicates anti-His tag (anti-mouse-IgG secondary) fluorescence and red coloring indicates anti- Gαs (anti-Rabbit IgG secondary) fluorescence. Middle right: preparative size-exclusion chromatography traces with the approximate location of the complex flanked by dashed lines. Right: Analytical size-exclusion chromatography traces of the pooled complex fractions that had been flash-frozen and thawed before injection. “Sol.” Indicates the solubilized insect cell fraction and “F.T.” indicates anti-FLAG column flow-through.

**Figure S6:**
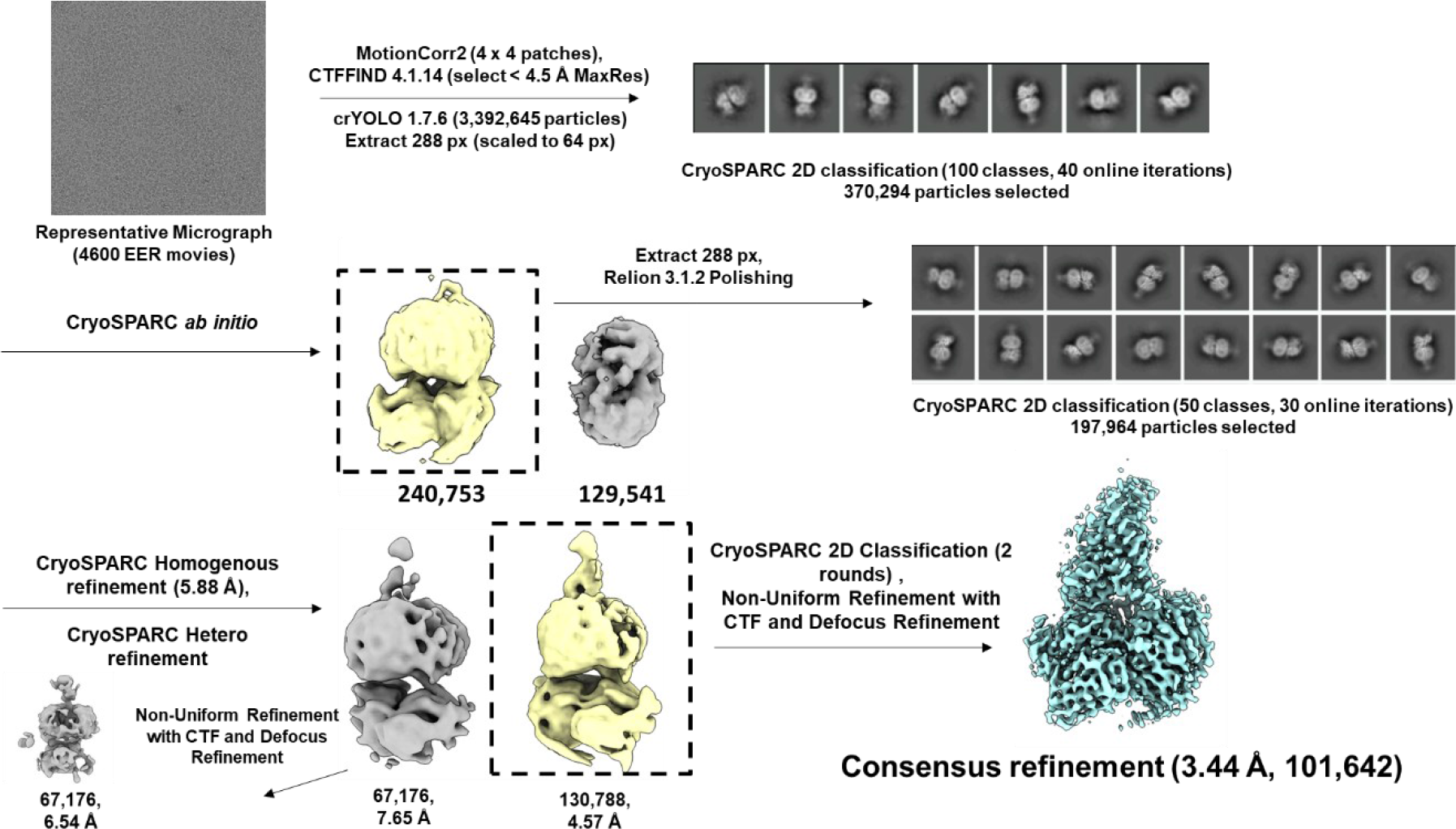
Overview of the cryo-EM data processing pipeline for the complex of peptide **1** bound to GLP- 1R/Gs. The field of view for the representative micrograph is 360 x 360 nm.

**Figure S7:**
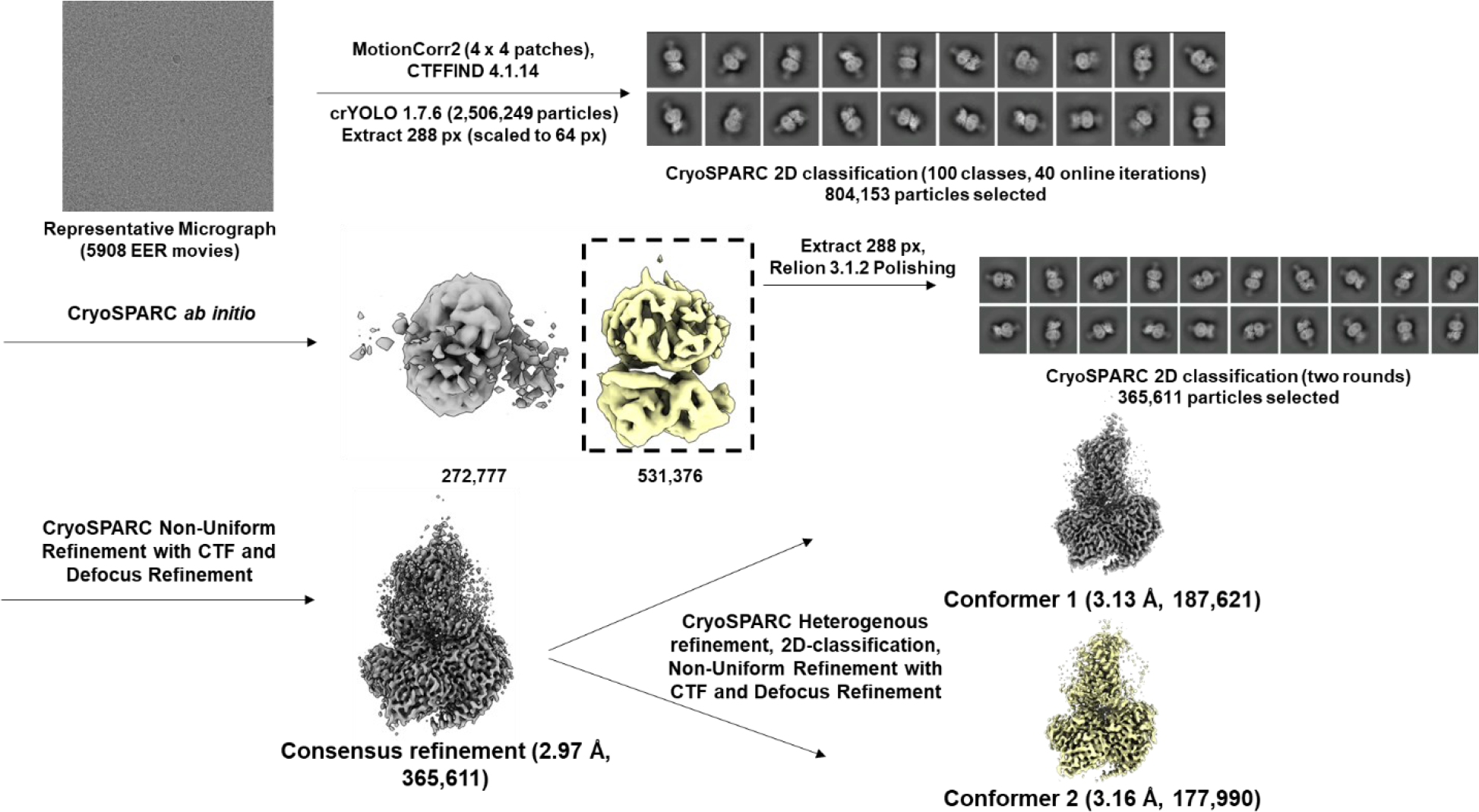
Overview of the cryo-EM data processing pipeline for the complex of peptide **2** bound to GLP- 1R/Gs. The field of view for the representative micrograph is 360 x 360 nm.

**Figure S8:**
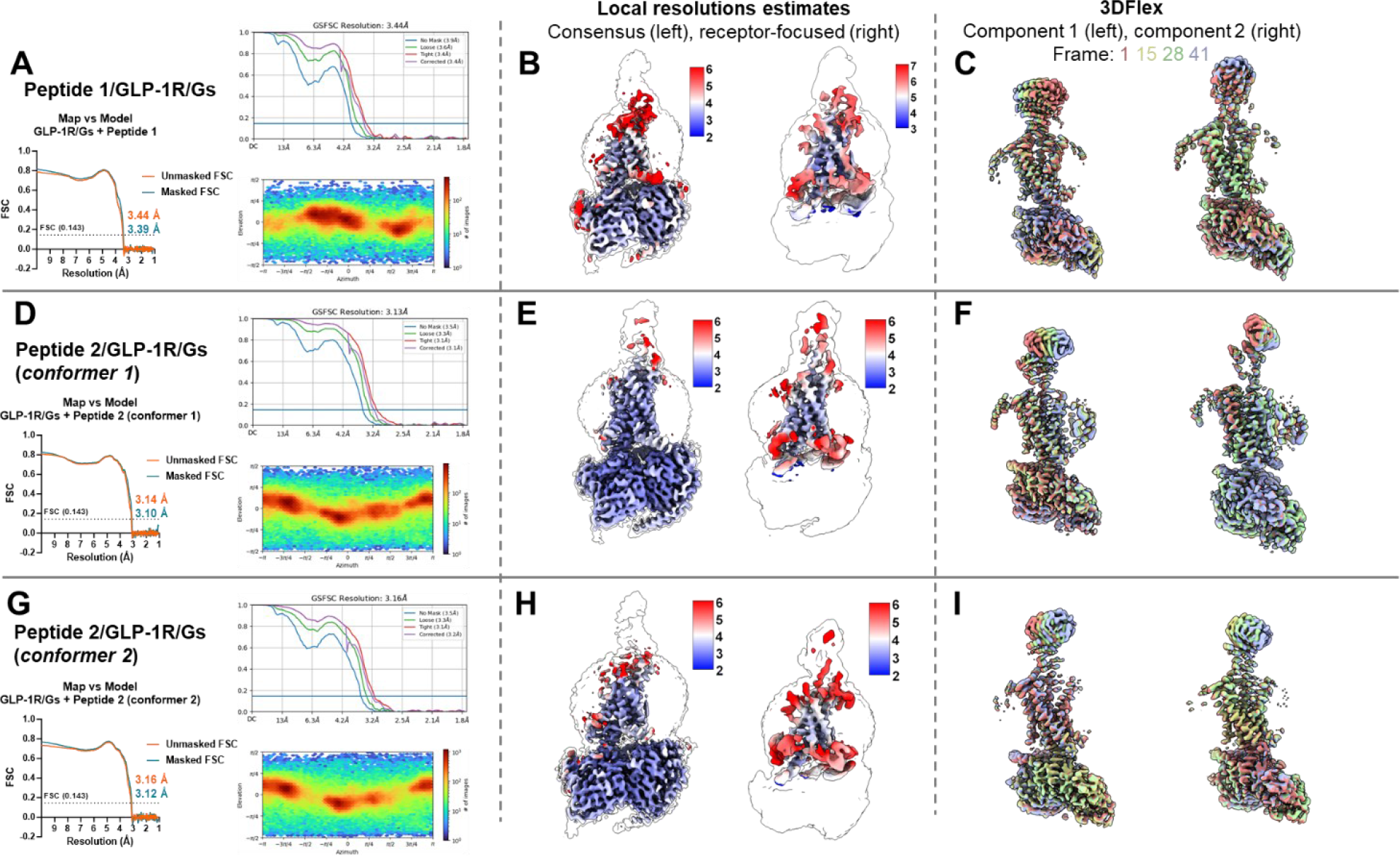
Cryo-EM reconstructions and resolution information for Peptide 1/GLP-1R/Gs (**A, B, C**), Peptide 2/GLP-1R/Gs (conformer 1) (**D,E,F**), and Peptide 2/GLP-1R/Gs (conformer 2) (**G,H,I**). For subfigures (**A,D,G**) Top: Gold standard Fourier shell correlation (FSC) curves with curves for various masks generated by Cryosparc 3.2. Resolutions were calculated using an FSC threshold of 0.143 (shown as a horizontal blue line). Bottom: Particle distribution heatmap. (**B,E,H**) Left: Estimated local resolution heatmap for the consensus refinement. Right: Estimated local resolution heatmap for the local refinement of the receptor and ECD. (**C,F,I**) CryoSPARC 3D Flex reconstructions showing representative frames (frame numbers 1, 15, 28 and 41 out of 41 total frames) of the 1^st^ and 2^nd^ latent components (left and right, respectively).

**Figure S9:**
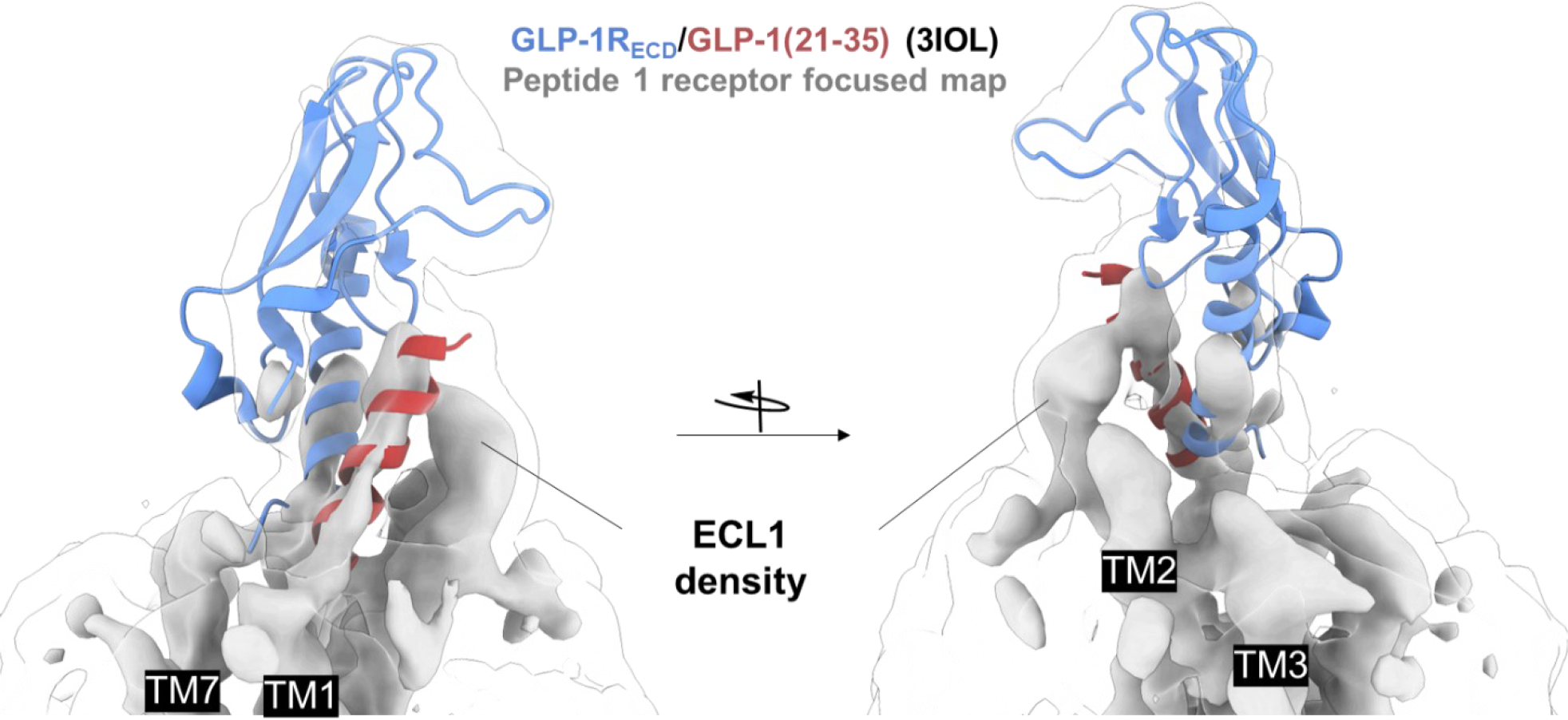
The crystal structure of GLP-1 bound to the GLP-1R extracellular domain (ECD, PDB: 3IOL) rigidly fit to the receptor-focused EM density map for peptide 1 bound to GLP-1R. The transparent map was contoured at low threshold, and the opaque map was contoured at lower threshold.

**Figure S10:**
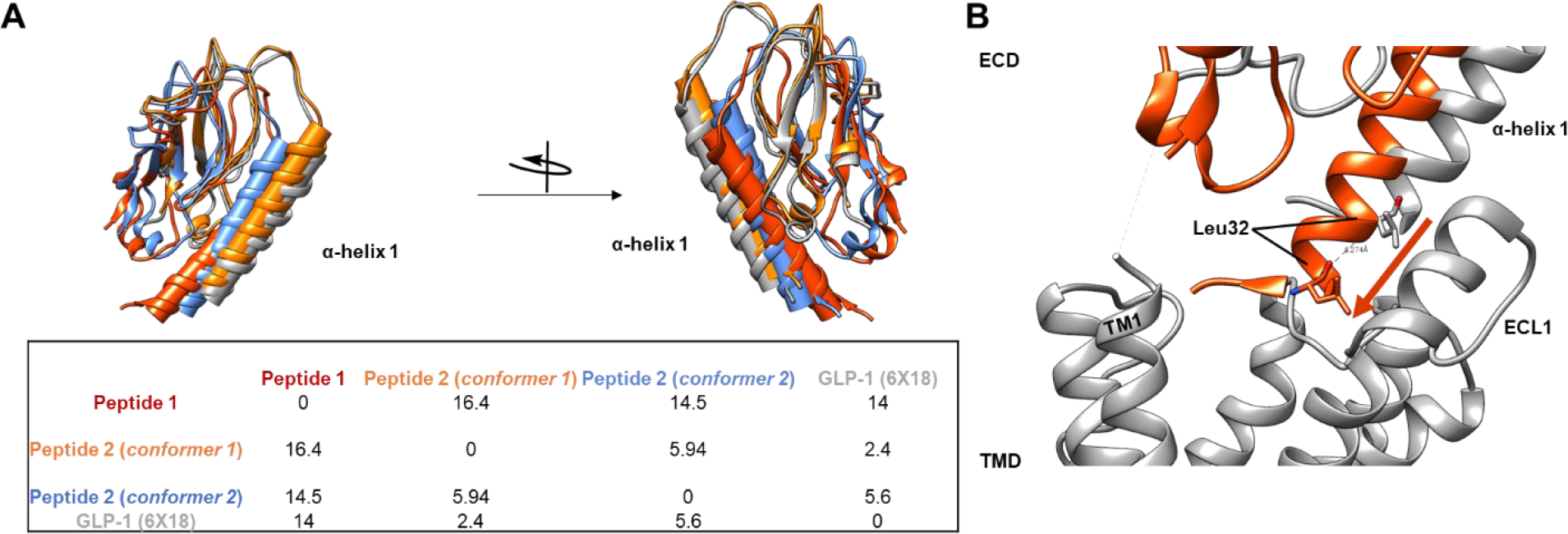
Comparison of receptor extracellular domain conformations inferred by cryo-EM. (A) Receptor-focused cryo-EM maps of complexes containing peptide **1** and **2** (conformers 1 and 2) were aligned, and a copy of the crystal structure of the GLP-1R extracellular domain (PDB: 3IOL) was rigidly fit into each. The full-length cryo-EM structure of GLP-1/GLP-1R (PDB: 6X18) was rigidly fit into the EM map of peptide **1** for reference. The inset table shows crossing angles for α-helix 1 (shown as cylinders in the figure) of the GLP-1R ECD in degrees. (B) Using the same model fits described above, the distance between the ECD α-helix 1 for GLP-1 (gray) and peptide **1** bound models (orange) was determined by the Cα atom of Leu32 in each model. Map fitting, angle, and distance calculations were performed using UCSF Chimera 1.14.

**Figure S11:**
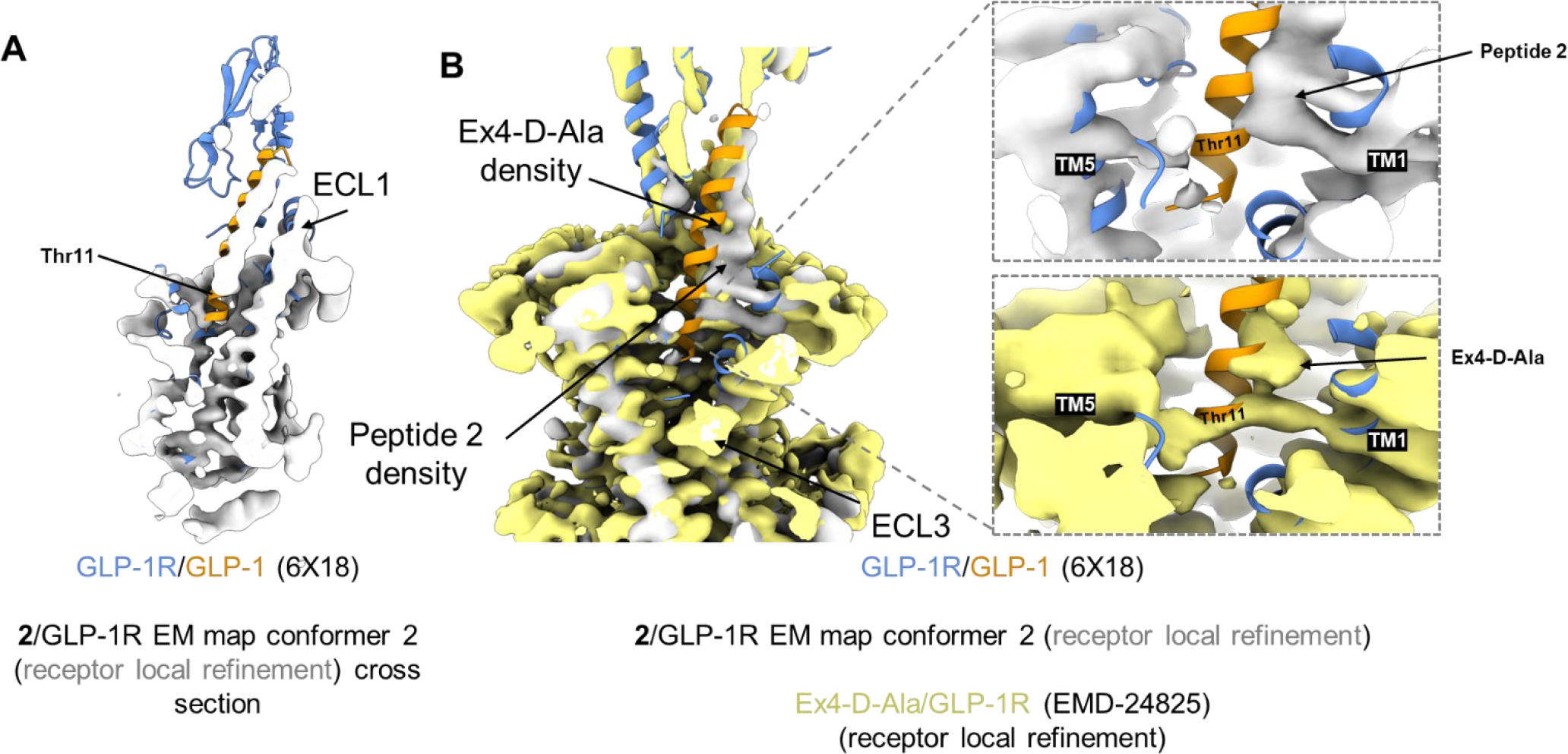
Peptide 2 bound to GLP-1R/Gs conformer 2. (A) Shown is a cutaway of the cryo-EM density for peptide 2 bound to GLP-1R/Gs (conformer 2, receptor-focused refinement). GLP-1 bound to GLP- 1R/Gs (PDB 6X18) was fit to the cryo-EM map, and it indicates that the density for peptide 2 (conformer 2) is absent near the N-terminus of the fitted GLP-1 peptide. (B) A comparison of the aligned cryo-EM density maps for peptide 2 (conformer 2), shown in gray, and Ex4-D-Ala (conformer 2) (EMD-24825), shown in gold. GLP-1 bound to GLP-1R/Gs is fitted to the map of peptide 2 for reference.

**Table S1:**
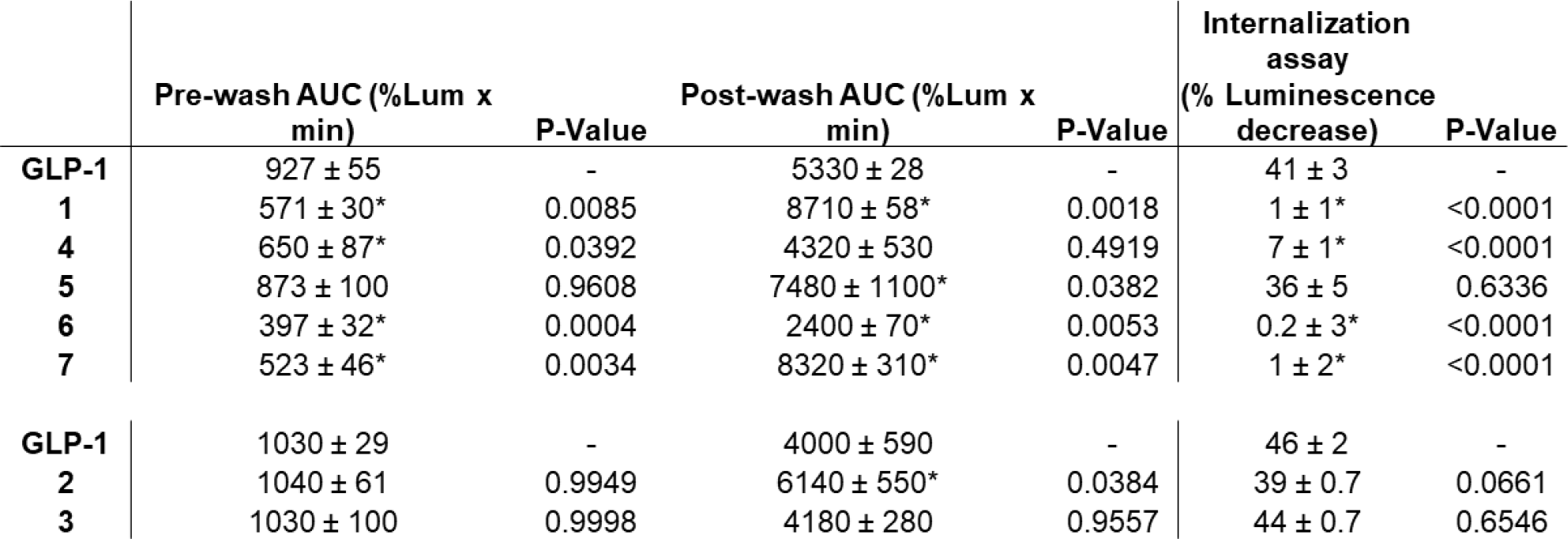
Tabulated results from washout and internalization assays where final data were not normalized to GLP-1. Experiments were performed in two batches of assays (each batch included GLP-1 as a control and either peptides 1,4-7 or peptides 2 and 3), which were analyzed independently by one-way ANOVA with Dunnett’s post-test compared to GLP-1 to generate P-values. * indicates P < 0.05. % Luminescence decrease was measured as the loss in luminescence relative to cells treated with vehicle. Larger losses in luminescence indicates more extensive nLuc-GLP-1R internalization. Uncertainties are shown as standard error of the mean. n = 3 for all experiments, except for the internalization assays with GLP-1, peptide 2 and peptide 3, where n =2.

**Table S2:**
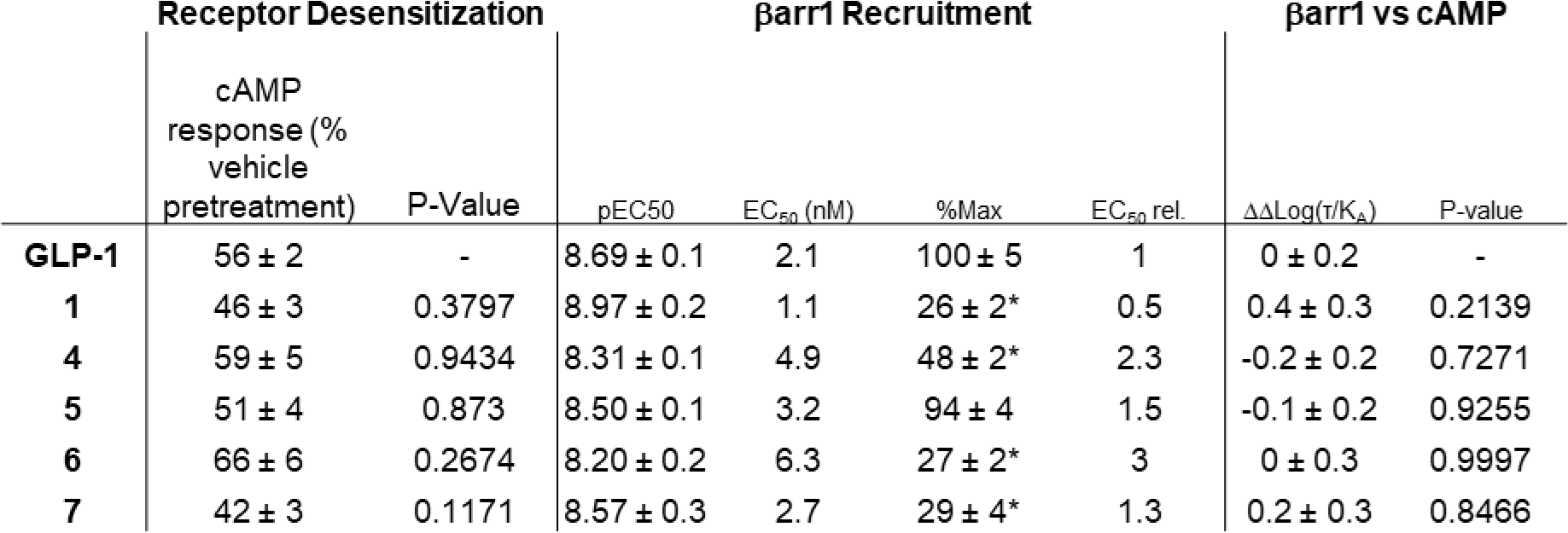
Best fit parameters from concentration-response experiments and tabulated results from experiments from Figures 2B and 2F. EC_50 rel._ indicates arrestin recruitment potency relative to GLP-1 by the quotient (analogue EC_50_)/(GLP-1 EC_50_). Bias factors (DDlog(t/K_A_)) were calculated in terms of b-arrestin-1 recruitment relative to cAMP production, and relative to GLP-1. Positive bias factor values indicate bias towards arrestin-recruitment and negative values indicate bias towards cAMP-production, relative to GLP-1. Uncertainties represent standard error of the mean except for bias factors where errors are expressed as standard deviation. n = 3 for arrestin recruitment assays and n = 4 for receptor desensitization assays. P-values were calculated by one-way ANOVA with Dunnett’s post-test compared to GLP-1. * indicates P < 0.05 compared to GLP-1.

| | Receptor Desensitization | | βarr1 Recruitment | | | | βarr1 vs cAMP | |
| --- | --- | --- | --- | --- | --- | --- | --- | --- |
| | cAMP response (% vehicle pretreatment) | P-Value | pEC<sub>50</sub> | EC<sub>50</sub> (nM) | %Max | EC<sub>50</sub> rel. | $\Delta\Delta\text{Log}(\tau/K_A)$ | P-value |
| **GLP-1** | 56 ± 2 | - | 8.69 ± 0.1 | 2.1 | 100 ± 5 | 1 | 0 ± 0.2 | - |
| **1** | 46 ± 3 | 0.3797 | 8.97 ± 0.2 | 1.1 | 26 ± 2* | 0.5 | 0.4 ± 0.3 | 0.2139 |
| **4** | 59 ± 5 | 0.9434 | 8.31 ± 0.1 | 4.9 | 48 ± 2* | 2.3 | -0.2 ± 0.2 | 0.7271 |
| **5** | 51 ± 4 | 0.873 | 8.50 ± 0.1 | 3.2 | 94 ± 4 | 1.5 | -0.1 ± 0.2 | 0.9255 |
| **6** | 66 ± 6 | 0.2674 | 8.20 ± 0.2 | 6.3 | 27 ± 2* | 3 | 0 ± 0.3 | 0.9997 |
| **7** | 42 ± 3 | 0.1171 | 8.57 ± 0.3 | 2.7 | 29 ± 4* | 1.3 | 0.2 ± 0.3 | 0.8466 |

**Table S3:** Best fit parameters from concentration-response experiments from Figure **S3B**. cAMP production (15 min) indicates fitted values from concentration-response curves generated from cAMP measurements ∼15 min post peptide addition. Uncertainties represent standard error of the mean. n = 3.

|  | cAMP Production |  |  |
| --- | --- | --- | --- |
|  | pEC<sub>50</sub> | EC<sub>50</sub><br>(nM) | % Max |
| **GLP-1-TMR** | 10.6 ± 0.08 | 0.024 | 98 ± 3 |
| **1-TMR** | 9.89 ± 0.05 | 0.13 | 97 ± 2 |
| **GLP-1** | 10.8 ± 0.1 | 0.015 | 100 ± 3 |
| **1** | 9.71 ± 0.05 | 0.2 | 97 ± 2 |

**Table S4:** Cryo-EM imaging conditions, refinement statistics, and model statistics. EER indicates electron event representation, a data format in which frames for dose-fractionation were not predetermined. * images were collected and processed with a pixel size of 0.8817 Å, but maps were changed to a pixel size of 0.878 Å after instrument calibration.

|  | *Peptide 1/GLP-1R/Gs* | *Peptide 2/GLP-1R/Gs* |  |  |
| --- | --- | --- | --- | --- |
| **Imaging** |  |  |  |  |
| Magnification | 120,000 | 120,000 |  |  |
| Voltage | 200 kV | 200 kV |  |  |
| Spot Size | 5 | 5 |  |  |
| Electron exposure ($e^-/\text{\AA}^2$) | 50 | 50 | | |
| Exposure time (s) | 7.33 | 7.39 |  |  |
| Movie frames | N/A (EER) | N/A (EER) |  |  |
| Pixel size ($\text{\AA}$) | 0.878* | 0.878* | | |
| AFIS | Yes | Yes |  |  |
| **Processing** | *Consensus map* | *Consensus map* | *Conformer 1* | *Conformer2* |
| Symmetry imposed | C1 | C1 | C1 | C1 |
| Initial particle images (no.) | 3,392,645 | 2,506,249 | 2,506,249 | 2,506,249 |
| Final particle images (no.) | 101,642 | 365,661 | 187,621 | 177,990 |
| Resolution ($\text{\AA}$) | 3.44 | 2.97 | 3.13 | 3.16 |
| FSC threshold | 0.143 | 0.143 | 0.143 | 0.143 |
| **Refinement** |  |  |  |  |
| Initial model used | *De novo* | *De novo* | *De novo* | *De novo* |
| B-factor | 92.2 | 105.6 | 89.9 | 81.6 |
| **Model composition** |  |  |  |  |
| Chains | 5 | - | 6 | 5 |
| Non-Hydrogen Atoms | 7639 | - | 8857 | 7640 |
| Protein residues | 1001 | - | 1153 | 1003 |
| Ligands | 0 | - | 1 | 0 |
| **RMSDs** |  |  |  |  |
| Bond length ($\text{\AA}$) | 0.005 | - | 0.006 | 0.005 |
| Bond angles ($^\circ$) | 0.953 | - | 0.966 | 0.965 |
| **Validation** |  |  |  |  |
| MolProbity score | 1.58 | - | 1.51 | 1.60 |
| Clashscore | 6.09 | - | 4.23 | 6.90 |
| Rotamer outliers (%) | 0 | - | 0.79 | 0.79 |
| **Ramachandran plot** |  |  |  |  |
| Favored (%) | 96.33 | - | 95.66 | 96.64 |
| Allowed (%) | 3.57 | - | 4.34 | 3.26 |
| Outliers (%) | 0.1 | - | 0 | 0.1 |

