## Supplemental Information for "Prolonged signaling of backbone-modified glucagon-like peptide-1 analogues with diverse receptor trafficking"

**Author affiliations:** [1]Department of Chemistry, University of Wisconsin − Madison, 1101 University Avenue, Madison, Wisconsin 53706, United States [2]Drug Discovery Biology Theme, Monash Institute of Pharmaceutical Sciences, Monash University, Parkville 3052, Victoria, Australia. [3]ARC Centre for Cryo-Electron Microscopy of Membrane Proteins, Monash Institute of Pharmaceutical Sciences, Monash University, Parkville 3052, Victoria, Australia

#### Supplementary figures and tables:

### Materials and Instrumentation:

#### A. Instrumentation:

Automated solid-phase peptide synthesis (SPPS) was performed with a CEM Liberty Blue instrument with an $HT_{12}$ resin-handling module. Manual, microwave assisted SPPS was performed with Torviq polypropylene syringes fitted with a porous polypropylene disk at the bottom using a CEM MARS microwave instrument. Preparative HPLC was performed using either a Shimadzu HPLC system (SCL-10VP system controller, LC-6AD pumps, SIL-10ADVP autosampler, SPD-10VP UV-vis detector, FRC-10A fraction collector) or a Waters HPLC system (Waters 2545 Binary Gradient module, Waters 2707 Autosampler, Waters 2998 Photodiode Array Detector, and Waters Fraction Collector III) . The HPLC systems were outfitted with a C18 column (either Agilent ZORBAX 21.2x250 mm, 7 µm or Waters XSelect CSH 19x250 mm, 5 µm). Matrix-assisted laser desorption ionization-time-of-flight (MALDI-TOF) mass spectrometry was performed by pipetting 1 µL of peptide mixture and 1 µL of CHCA (saturated solution in 1:1 acetonitrile:water) onto an appropriate target, allowing the liquid to evaporate completely under ambient conditions, and acquiring spectra on a Bruker microflex™ LRF. High resolution mass measurements were performed with a Bruker impact II ESI-qTOF system. Liquid-Chromatography-Mass Spectrometry (LCMS) was performed on a Waters Acquity Arc system outfitted with a Waters XBridge C18 1.2 x 50 mm (3.5 µm) column. UV-Vis measurements of peptides and plasmids were performed with with either a ThermoFisher NanoDrop 2000 or a NanoDrop One C microvolume spectrophotometer. Luminescence and bioluminescence resonance energy transfer (BRET) were measured with a BioTek Synergy 2 plate reader.

#### B. Instrument acknowledgements:

The purchase of the Bruker microflex™ LRF in 2015 was funded by the Bender gift to the Department of Chemistry.

#### C. Table of Materials:

| Item/Reagent | Vendor | Item # | Notes |
|---|---|---|---|
| **α-N-Fmoc-amino acids** | Chem Impex | Various | |
| **Fmoc-ACPC** | Chem Impex | 15073 | |
| **Fmoc-APC(Boc)-OH** | Synthesized in house | - | See LePlae et al.[1] for synthetic route |
| **6-Carboxy-tetramethylrhodamine** | Chem Impex | 16901 | |
| **NovaPEG Rink Amide** | Milipore-Sigma | 8550470005 | |
| **SPPS Reaction Vessel** | Torviq | SPE-0500 | (5mL) |
| **O-(7-Azabenzotriazol-1-yl)-N,N,N',N'-tetramethyluronium hexafluorophosphate (HATU)** | Chem Impex | 12881 | |
| **N,N'-Diisopropylcarbodiimide (DIC)** | Chem Impex | 00110 | |
| **Ethyl (hydroxyimino)cyanoacetate (Oxyma)** | Chem Impex | 26426 | |
| **α-Cyano-4-hydroxycinnamic acid (CHCA)** | Sigma-Aldrich | 70990 | |
| **Diisopropylethylamine (DIEA)** | Sigma-Aldrich | 496219 | Biotech. Grade |
| ***N*, *N* Dimethylformamide (DMF)** | Sigma-Aldrich | 319937 | ACS Reagent |

| | | | |
|---|---|---|---|
| ***N*, *N* Dimethylformamide (DMF)** | Sigma-Aldrich | 494488 | Biotech. Grade |
| **Piperidine** | Sigma-Aldrich | 104094 | |
| **Piperazine** | Sigma-Aldrich | P45907 | |
| **Trifluoroacetic acid** | Sigma-Aldrich | T6508 | ReagentPlus, 99% |
| **1,2 Ethanedithiol** | Sigma-Aldrich | 2390 | |
| **Thioanisole** | Sigma-Aldrich | T28002 | |
| **D-Luciferin** | GoldBio | LUCK-100 | (Potassium Salt) |
| **Coelenterazine 400 a** | GoldBio | C-320-10 | also known as DeepBlueC™ |
| **H-Coelenterazine** | Nanolight | 301 | |
| **FuGENE® HD** | Promega | E2311 | |
| **Nano-Glo® Luciferase Assay System** | Promega | N1110 | |
| **pCMV6-XL5 [hGLP1-R]** | Origene | SC124060 | |
| **nLuc-GLP-1R-His-FLAG** | In house | Available at Addgene (# 124831) | See Cary et al.[2] |
| **GLP-1R-Rluc8** | In house | - | See Hager et al.[3] |
| **pEGFP-N2 [GLP-1R-eGFP]** | - | - | Kind gift from Alessandro Bisello. See Syme et al.[4] |
| **pCMV6-XL5 [GFP²ß-Arrestin-1]** | In house | - | See Hager et al.[3] |
| **GFP²ß-Arrestin-2(R393E, R395E)** | In house | - | See Hager et al.[3] |
| **GRK5** | - | - | Kind gift from Rasmus Jorgensen and Jakob Lerche Hansen (Novo Nordisk) |
| **HEK293FT Cells** | ThermoFisher | R70007 | |
| **HEK293-GS22 Cells** | Promega | E1261 | Kind Gift from Thomas Gardella |
| **DMEM** | Gibco (ThermoFisher) | 11965092 | +High glucose, L-Gln, Phenol Red |
| **Opti-MEM** | Gibco (ThermoFisher) | 31985070 | With L-Gln |
| **McCoy's 5A Medium Modified** | Gibco (ThermoFisher) | 12330031 | +High glucose, L-Gln, Bacto-peptone, Phenol Red, Hepes |
| **DPBS** | Sigma-Aldrich | D8537 | |
| **$CO_2$ Independent Medium** | Gibco (ThermoFisher) | 18045088 | |
| **Bovine Albumin Fraction V (7.5% solution)** | Gibco (ThermoFisher) | 15260037 | |
| **DPBS (+Glucose)** | Gibco (ThermoFisher) | 14287080 | +calcium, magnesium, glucose, pyruvate |
| **100X L-Glutamine** | Gibco (ThermoFisher) | 25030081 | |
| **100X Sodium Pyruvate** | Gibco (ThermoFisher) | 11360070 | |
| **100X MEM NEAA** | Gibco (ThermoFisher) | 11140050 | |
| **Hyclone™ 100X Penicillin-Streptomycin** | GE Healthcare Life Sciences | SV30010 | |
| **Fetal Bovine Serum** | Gibco | 10082147 | Heat Inactivated |
| **10 cm, tissue-treated dishes** | Corning | 353003 | |
| **75 cm² vented, tissue-treated flasks** | Corning | 430641U | |
| **0.05% Trypsin-EDTA** | Gibco (ThermoFisher) | 25300054 | With Phenol Red |

| | | | |
|---|---|---|---|
| **96-well plates (white bottom)** | Corning | 3917 | TC-treated, white, sterile |
| **EndoFree Plasmid Maxi Kit** | Qiagen | 12362 | |
| **Human GLP-1(7-36)-NH$_2$** | Prospec | HOR-284 | |
| **Marimastat** | Alfa Aesar (ThermoFisher) | J67288 | |
| **Glass-bottom dishes** | MatTek | P35G-1.5-10-C | 35 mm Dish \| No. 1.5 Coverslip \| 10 mm Glass Diameter \| Uncoated |
| **Poly-L-Lysine solution** | Sigma | P4832 | Mol weight 150-300k, 0.01% |
| **16% Paraformaldehyde (formaldehyde) aqueous solution** | Electron Microscopy Sciences | 15700 | |
| **Hoechst 33342** | ThermoFisher | H3570 | Trihydrochloride, Trihydrate - 10 mg/mL Solution in Water |
| **UltrAuFoil R1.2/1.3 grids** | Electron Microscopy Sciences | Q350AR13A | |
| **Exendin (9-39)-NH$_2$** | Genscript | RP10872 | |

### Cell culture:

HEK293-GS22 cells were maintained in 75 cm$^2$, culture-treated, vented flasks (Corning) at in a humidified atmosphere at 37°C with 5% $CO_2$. Cell medium was 0.22 µm-filtered DMEM supplemented with 10% (v/v) FBS and penicillin/streptomycin. Cells were subcultured every 4-5 days at confluency.

HEK293FT cells were maintained under similar conditions except their medium was supplemented with 100-fold diluted 100X L-Glutamine, 100X Sodium Pyruvate and, 100X MEM NEAA to working concentrations of 1X. After receiving cells, cultures were not tested for mycoplasma contamination.

#### Transient transfections:

Except for the arrestin recruitment assays, all assays and microscopy were conducted with transiently transfected HEK293 cells stably expressing the Glosensor-22F (Promega) luminescent cAMP-sensing protein. Cells were grown to confluence in 75 cm$^2$ flasks, harvested, and then 1/3 of the collected cells were plated onto a 10 cm tissue-treated dish with 10 mL 10% FBS in DMEM without penicillin/streptomycin. The cells were then incubated at 37°C with 5% $CO_2$ overnight. After the overnight incubation, the medium was aspirated, 4.5 mL McCoy's 5A modified medium with 10% FBS was added, and the cells were incubated at 37°C with 5% $CO_2$. During this incubation, plasmid (as a solution in endotoxin-free TE buffer (Qiagen)) and FuGENE HD transfection reagent were added to 1 mL of opti-MEM. After 20 min, 4.5 mL of DMEM with 10% FBS was added to the cells, and 1 mL of the transfection mixture was gently pipetted into the medium. The cells were then returned for incubator overnight.

### Luminescence and BRET assays:

#### Single-timepoint cAMP assays:

The protocol was adapted from Hager et al.[3] and Binkowski et al.[5] HEK293 cells stably expressing the Glosensor-22F (Promega) luminescent cAMP-sensing protein were transiently transfected with either 1.2

μg of nluc-GLP-1R plasmid and 5 μL FuGENE HD transfection reagent or with 10 μg of hGLP-1R plasmid and 30 μL FuGENE HD transfection reagent. The next day, cells were washed with DPBS, harvested with 0.05% Trypsin-EDTA, and resuspended into 4 mL 10% FBS in DMEM without penicillin/streptomycin. The cell suspension was diluted to approximately 500,000 cells/mL in medium, and 100 μL of cell suspension was pipetted into each well (providing ~50,000 cells/well) of a white-bottom, white-walled, 96-well plate. The cells in the 96-well plate were incubated at 37°C with 5% $CO_2$ overnight. After 24 h, the medium was removed by inverting and gently flicking the plate. DPBS with D-luciferin (500 μM) was quickly added to the plate (90 μL/well). The plate containing cells was allowed to sit for approximately 20 min at room temperature before addition of peptide (as 10 μL/well diluted in DPBS). We found it to be important to change pipette tips between serial dilutions for reproducible results. The plate was then transferred to a BioTek Synergy 2 plate reader with no optical filter ("hole"), 1 mm vertical probe offset, and read with a sensitivity value of 200. Curves were generated from luminescence values observed between 10 and 20 min.

Experiments were conducted with $n = 3$ biological replicates. Reported $EC_{50}$ and %Max values were a result of normalizing, averaging, and then fitting data to three-parameter sigmoidal curves in GraphPad Prism 6. The bottom of the curves was constrained to 0%. Normalization was performed with 100% representing the top of GLP-1's curve for individual experiments and 0% representing the luminescence value in absence of peptide. Reported errors in maximal response and $EC_{50}$ were estimated from the fitted data using GraphPad Prism.

#### cAMP washout assays:

The protocol was adapted from Liu et al.[6] HEK293 cells stably expressing the Glosensor-22F (Promega) luminescent cAMP-sensing protein were transiently transfected with 10 μg of hGLP-1R plasmid and 30 μL FuGENE HD transfection reagent. The next day, cells were washed with DPBS, harvested with 0.05% Trypsin-EDTA, and resuspended into 4 mL 10% FBS in DMEM without penicillin/streptomycin. The cell suspension was diluted to approximately 500,000 cells/mL in medium, and 100 μL of cell suspension was pipetted into each well (providing ~50,000 cells/well) of a white-bottom, white-walled, 96-well plate. The cells in the 96-well plate were incubated at 37°C with 5% $CO_2$ overnight. After 24 h, the medium was removed by inverting and gently flicking the plate. $CO_2$ independent medium with 0.1% BSA and D-luciferin (500 μM) was quickly added to the plate (90 μL/well). The plate containing cells was allowed to sit for approximately 20 min at room temperature before addition of peptide (as 10 μL/well diluted in DPBS). The plate was then transferred to a BioTek Synergy 2 plate reader with no optical filter ("hole"), 1 mm vertical probe offset, and read with a sensitivity value of 200. After 15 minutes of luminescence reading, the plate was washed twice with $CO_2$ independent medium (100 μL/well), and fresh media ($CO_2$ independent medium with 0.1% BSA and D-luciferin (500 μM)) with or without exendin-9-39 (100 nM) was added (100 μL/well). Luminescence was then monitored over four hours.

#### cAMP desensitization assays:

Desensitization assays were performed in an identical manner to washout assays with 10 nM agonist except D-luciferin and exendin-9-39 were not included before or during the washout. After the washout and a four-hour incubation at 37°C, D-luciferin and GLP-1 in solutions of DPBS were added to the well to final concentrations of 500 μM and 100 nM respectively. The plate was then transferred to a BioTek Synergy 2 plate reader with no optical filter ("hole"), 1 mm vertical probe offset, and read with a sensitivity value of 200.

#### Bioluminescence resonance energy transfer (BRET) arrestin recruitment assays:

The protocol was adapted from Hager et al.[3] and Jorgensen et al.[9] HEK293FT cells were grown confluence, harvested, and then 1/3 of the collected cells were plated onto a 10 cm tissue-treated dish with 10 mL 10% FBS in DMEM (with NEAA, L-Glutamine, and Sodium pyruvate, see above) without penicillin/streptomycin. The cells were then incubated at 37°C with 5% $CO_2$ overnight. After 24 h, a transfection mixture was made with 1:1 polyethylenimine (PEI, 1 mg/mL in water, pH 7.0):DNA in 1 mL opti-MEM. $GFP^2$-ß-arrestin-1 (14 µg) along with GRK5 (250 ng) and GLP-1R-Rluc8 (250 ng) was added to the PEI/Opti-MEM mixture, and the resulting transfection mixture was incubated for 20 min at room temperature. The cell medium was then aspirated, replaced with 4.5 mL of DMEM without FBS, and the transfection mixture was gently pipetted into the medium. Six hours after transfection (with incubation at 37°C with 5% $CO_2$), 4.5 mL of DMEM supplemented with 20% FBS was added to the dish. Twenty-four hours after transfection (with incubation at 37°C with 5% $CO_2$), cells were washed with DPBS, harvested with 0.05% Trypsin-EDTA, and resuspended into 4 mL 10% FBS in DMEM (with NEAA, L-Glutamine, and Sodium pyruvate, see above). The cell suspension was diluted to approximately 1,000,000 cells/mL in medium, and 100 µL of cell suspension was pipetted to each well (providing ~100,000 cells/well) of a white-bottom, white-walled, 96-well plate. The cells in the 96-well plate were incubated at 37°C with 5% $CO_2$ overnight.

Twenty-four hours after adding cells to the 96-well plate, the medium was removed by pipette, the cells were washed twice with DPBS (with glucose, 100 µL/well), and 100µL of DPBS (with glucose) was added to each well. The cells were incubated at 37°C with 5% $CO_2$ for 45 min to1 h before addition of peptide (as 10 µL dilutions in DPBS). After addition of peptides, the cells were allowed to sit at room temperature for 20 min before addition of the Rluc8 substrate, DeepBlueC (10 µL/well of 60 µM DeepBlueC in 2:1 DPBS:ethanol). The 96-well plate was transferred to a BioTek Synergy 2 plate reader with 400 nm (20 nm bandwidth) and 528 nm (30 nm bandwidth) optical filters, 1 mm vertical probe offset, 1-second integration time and read at maximum (200) sensitivity. Concentration-response curves were generated with $I_{528nm}/I_{400nm}$ values taken between 15 and 45 min after the initial read, as signal variability was found to be relatively high at earlier timepoints.

Experiments were repeated with n = 3 biological replicates. Each experiment consists of at least 7 different concentrations of peptide, with solutions prepared via serial dilution of a stock solution of each peptide prepared from 1 mM DMSO stock. Reported $EC_{50}$ and %Max values were a result of normalizing, averaging, and fitting data to three-parameter sigmoidal curves in GraphPad Prism 6. The bottom of the curves was constrained to 0%. Normalization was performed with 100% representing the top of GLP-1's curve for individual experiments and 0% representing the bottom value if the curves were fit with raw data and constrained to have a shared minimum. Reported errors in maximal response and $EC_{50}$ were estimated from the fitted data using GraphPad Prism.

##### Receptor internalization assays:

HEK293 cells stably expressing the Glosensor-22F (Promega) luminescent cAMP-sensing protein were transiently transfected with 1.2 µg of nluc-GLP-1R plasmid and 5 µL FuGENE HD transfection reagent. The next day, cells were washed with DPBS, harvested with 0.05% Trypsin-EDTA, and resuspended into 4 mL 10% FBS in DMEM without penicillin/streptomycin. The cell suspension was diluted to approximately 500,000 cells/mL in medium, and 100 µL of cell suspension was pipetted into each well (providing ~50,000 cells/well) of a white-bottom, white-walled, 96-well plate. The cells in the 96-well plate were incubated at 37°C with 5% $CO_2$ overnight. The next day, the media was removed and replaced with $CO_2$ independent medium supplemented with 0.1% BSA and 1 µM marimastat. Peptide solutions were added as 10 µL aliquots in DPBS, and the cells were incubated at 37°C. After 30 min., a solution of H-

Coelenterazine (2 mL DPBS + 100 μL of 1.4 mM H-coelenterazine in ethanol) was prepared and added to the 96-well plate (10 μL solution/well). The plate was then transferred to a BioTek Synergy 2 plate reader and read with a 1 s integration time over the course of 1 h with 460 nm (40 nm bandwidth).

The maximal luminescence values for each well were selected after 5 minutes post-addition of H-coelentrazine. Wells treated with vehicle were averaged, and that mean value was normalized to 0% decrease in luminescence.

##### Confocal microscopy:

HEK293 cells stably expressing the Glosensor-22F (Promega) luminescent cAMP-sensing protein were transiently transfected with 5 μg of GLP-1R-eGFP plasmid and 15 μL FuGENE HD transfection reagent. The next day, poly-L-lysine solution (Sigma, 100 μL) was applied to glass bottom dishes (MatTek) for 15 min., the dishes were washed twice with DPBS (100 μL), and the dishes were allowed to dry for 1 h. The transfected cells were washed with DPBS, harvested with 0.05% Trypsin-EDTA, and resuspended into 10 mL 10% FBS in DMEM without penicillin/streptomycin. Cell suspension (2 mL) was added to each poly-L-lysine treated dish, and the dishes were then returned to the incubator. The next day, the media was removed from the dishes and TMR-labeled peptide solution (200 μL/dish, DPBS with glucose + 0.1% BSA) was added. The cells were incubated at 37°C for 30 min. Then, the peptide solution was aspirated, and the cells were washed with DPBS with glucose + 0.1% BSA (1 mL). The cells were fixed with 4% paraformaldehyde in DPBS, washed with DPBS, and excess aldehyde was quenched with a glycine wash. 1 mL of Hoescht 33342 (1:2000 in DPBS) solution was added for 5 min, and then the cells were then washed twice with DPBS. The cells were imaged with a Nikon A1R-SI+ laser-scanner confocal microscope system equipped with a 60X oil-immersion objective lens.

#### Nonlinear fitting:

Data were processed in Microsoft Excel and GraphPad Prism 6 software packages. Concentration-response data was normalized for each experiment in Prism (*vide supra* for normalization parameters). Averages and sample standard deviations were calculated in Excel. This processed data was returned to Prism for curve fitting. $EC_{50}$ and maximal-response values, as well as curves, shown in figures and tables are from 3-parameter sigmoidal fits.

We applied the Black-Leff operation model to quantify the relative efficacy of each peptide in activating the GLP-1R. Each set of peptides was fit to the equation below (Eq 1.) in GraphPad Prism 6, and transduction coefficients, or $\log(\tau/K_A)$ values, were extracted (as mean and SEM). $E$ represents the system response given a set of parameters. $E_{max}$ is the maximal response, $[A]$ is the concentration of the agonist, $K_A$ is the association constant of the agonist for the receptor, and $\tau$ is the receptor density $[R_t]$ divided by the intrinsic agonist efficacy $K_E$. Taking the difference of transduction coefficients between an agonist and the reference peptide (GLP-1(7-36)-$NH_2$) provides the normalized transduction coefficient, $\Delta\log(\tau/K_A)$. A bias-factor, or $\Delta\Delta\log(\tau/K_A)$ value, is calculated by determining the difference between the normalized transduction coefficients for two signaling pathways.

(Eq. 1)
$$E = \frac{E_{max}\tau[A]}{[A](1+\tau)+K_A}$$

Standard deviations for $\Delta\Delta\log(\tau/K_A)$ values were estimated by propagating extracted standard-errors for $\log(\tau/K_A)$ using equation 2. Where $\sigma_{prop}$. represents propagated standard deviation, and $SE^2_{ref1}$ and $SE^2_{ref2}$ represent the standard errors of the reference ligand (GLP-1) in the first and second pathway being compared, respectively. $SE^2_{lig1}$ and $SE^2_{lig2}$ represent the standard errors of the test ligand in the first and second pathway being compared, respectively.

(Eq. 2) $$\sigma_{prop.} = \sqrt{SE^2_{ref1} + SE^2_{ref2} + SE^2_{lig1} + SE^2_{lig2}}$$

To determine whether a $\Delta\Delta\log(\tau/K_A)$ value was statistically significant, $\Delta\Delta\log(\tau/K_A)$ values were compared using one-way analysis of variance (ANOVA) with Bonferroni's post-test.

**GLP-1R/Gs complex formation:** GLP-1R/Gs/Gβ1/Gγ2/Nanobody35 complexes were prepared as previously described with minor modifications.[2] 5 µM peptide was used during the initial complex formation step and 2 µM peptide was included in buffers for all other steps. Size-exclusion chromatography was performed with a flow rate at 0.7 mL/min.

**Cryo-EM:** UltrAuFoil R1.2/1.3 grids were glow discharged with a Quorum GloQube® Plus Glow Discharge System. Freshly thawed GLP-1R/Gs samples (3 µL, 4.7 or 4.3 mg/mL for samples including peptide **1** and peptide **2**, respectively) were pipetted onto grids in a ThermoFisher VitroBot Mark IV (4°C, 100% humidity), the grids were blotted (blot force 18, blot time 8 s), and samples were vitrified in liquid ethane.

The grids were clipped and loaded into a ThermoFisher Scientific Glacios transmission electron microscope outfitted with a Falcon 4 direct-electron detector (4096 x 4096 pixels). The microscope was operated at 200 kV and 120,000x indicated magnification in nanoprobe mode using spot size 5, a 50 µm C2 aperture, and a 100 µm objective aperture. Dosage was set to 50 $e^-/Å^2$, and exposure times were 7.33 s and 7.39 s for samples containing peptide **1** and peptide **2**, respectively. Automated data acquisition was performed with EPU 2.12.0.2771. Data were collected using 21-hole, aberration-free image shift (AFIS) and saved in electron-event representation (EER) format. For peptide **1** bound to GLP-1R/Gs, 4600 EER movies were collected, and for peptide **2** bound to GLP-1R/Gs, 5908 EER movies were collected. At the time of data acquisition, the physical pixel size was assigned as 0.8817 Å, but the pixel size was later recalibrated to 0.878 Å.

**Cryo-EM data processing:**

EER files were pre-processed using the EPU_Group_AFIS.py script (https://github.com/DustinMorado/EPU_group_AFIS) for import into RELION 3.1.2. MotionCor2 as implemented in RELION 3.1.2 was used for patch motion correction (4 X 4) with 37 EER fractionations. Contrast transfer function (CTF) estimation was performed with CTFFind 4.1.14. Particle picking was performed using CrYOLO 1.7.6. Scaled particles (288-pixel box size binned to 64-pixel box size) were extracted and imported into cryoSPARC 3.2.0 for one round of 2D classification followed by *ab initio* reconstruction with two classes. Selected particles were reextracted at full resolution, refined, and subjected to Bayesian polishing with RELION 3.1.2. Polished particles were imported into cryoSPARC 3.2.0 for additional rounds of 2D classification, heterogenous refinement, and finally non-uniform refinement with CTF and defocus refinement. A mask excluding density for the G protein and micelle was used to perform local refinement of the receptor and ECD density with rotation and shift search extents of 10 to 15° and 5 to 10 Å, respectively. 3D-variability was performed with cryoSPARC 3.2.0 using a filter resolution of 4 Å.

CryoSPARC 3DFlex was performed with default parameters (training box size of 128 px, 20 tetrahedral cells, two latent dimensions) using CryoSPARC 4.1.2. In addition to sharpened maps generated by cryoSPARC, post-processed maps were generated using DeepEMhancer using both half-maps.[10]

**Modeling:**

For modeling, maps were rescaled to a pixel size of 0.878 Å/px using UCSF Chimera, reflecting an microscope pixel size recalibration. We deposited rescaled maps, as well as the original (0.8817 Å/px) maps, half-maps, and FSC curves, in the Electron Microscopy Data Bank. The reported structure of GLP-1 bound to GLP-1R (PDB 6X18) was flexibly fit into the cryo-EM maps using ISOLDE 1.3 as implemented in UCSF ChimeraX 1.3. Real-space refinement was performed using Phenix 1.19.2 and WinCoot 0.9.6. For the ligand ACPC (XCP), restraint files were prepared using eLBOW as implemented in Phenix 1.19.2. Link files for XCP were prepared using JLigand (as part of CCP4i 7.1.017). Real-space validation was performed using Phenix 1.19.2.

#### Peptide Synthesis:

##### A. Peptide synthesis:

Polypeptides were generated by standard Fmoc-based solid-phase synthesis using a combination of automated and manual microwave-assisted methods.

For automated synthesis, NovaPEG Rink amide resin was swollen in DMF for 15 min and transferred to a Liberty blue automated peptide synthesis instrument. Couplings were performed with 2.5 mL of 0.2 M Fmoc-protected amino acid in DMF, 0.5 mL of 1 M oxyma, and 1 mL 1 M DIC with microwave irradiation at 75 °C for 3 min. Deprotections were performed with 20% piperidine in DMF with 0.1 M oxyma at 90 °C for 2 min.

Manual synthesis was performed in Torviq polypropylene syringes along with a Teflon coated magnetic stir-bar to complete the synthesis by manual, microwave-assisted methods. Fmoc-protected amino acids (100 µmol) were dissolved in 1 mL DMF (biotech. grade) with 37.5 mg/mL HATU. DIEA (35 µL) was added to this mixture, and the mixture was vortexed briefly and then incubated at room temperature for 2-3 minutes before addition to the resin/deprotected resin-bound polypeptide. Coupling was then performed with stirring and microwave irradiation to 70°C for either 4 min or 12 min. (We recommend that the first coupling, ß-residues, proline residues, arginine residues, and ß-branched residues are coupled for longer than 4 min under these conditions). Histidine residues were coupling for 12 min at 50 °C. Deprotections were performed by adding 4 mL 20% (v/v) piperidine in DMF with stirring and microwave irradiation to 80°C for 2 min. Resin was washed with at least 4 mL DMF (4x) after coupling and deprotection steps.

For tetramethylrhodamine labeled analogues, Nα-Fmoc-Nε-allyloxycarbonyl-L-lysine was incorporated into the polypeptide chain by standard microwave-assisted, Fmoc-based solid-phase peptide synthesis. After completing coupling steps for the entire peptide chain, the resin (25 μmol, NovaPEG Rink Amide) was washed with dichloromethane (3x, 4 mL). To achieve selective sidechain deprotection of the lysine protected with allyloxycarbonyl, a solution of $PhSiH_3$ (54.1 mg, 500 μmol) and $Pd(PPh_3)_4$ (7.2 mg, 6.25 μmol) in 1,2 dicholoroethane (1 mL) was freshly prepared and applied to the polypeptide-bound resin. The resulting mixture was stirred with microwave irradiation in open air at 35° C for 2 min. The resin was then washed with dichloromethane (4x, 4 mL). This deprotection procedure was performed twice. The resin was then washed with DMF and treated with a solution of 6-carboxytetramethyl rhodamine (21.8 mg, 50 μmol), HATU (18.8 mg, 49 μmol), and DIEA (17.5 μL) at room temperature with stirring for 24 h.

Cleavage of polypeptides from resin was performed with 3 mL peptide of 92.5% trifluoracetic acid (TFA), 5% thioanisole, and 2.5% 1,2-ethanedithiol (v/v) at room temperature with either stirring or rocking for 4 h. TFA was blown off with a stream of $N_2$, and the crude peptide was precipitated with addition of 30 mL of cold diethyl ether. Crude peptide was then pelleted by centrifugation at 3.5k RPM for 5 min. The supernatant was poured off, and the crude peptide pellet was dried under a $N_2$ stream.

For HPLC purification, the crude peptide pellet was then dissolved in either dimethyl sulfoxide (DMSO, 1-2 mL) or 1:1 acetonitrile: $H_2O$. 100-150 μL of peptide mixture was injected on the HPLC column and eluted with 12-14 mL/min of a 10-60% acetonitrile (with 0.1% TFA v/v) gradient in filtered Nanopure water (with 0.1% TFA v/v) over 60 minutes. Fractions were collected with a UV 220 nm-triggered automated fraction collector.

Peptide fractions were pooled, frozen over dry ice, and lyophilized. Lyophilized peptides were then dissolved in 1-2 mL of water with minimal acetonitrile to promote solubility of particularly hydrophobic peptides. Purity of the peptide was then determined by HPLC monitoring at 220 nm, and concentration was determined by UV-Vis absorption at 280 nm. Molar extinction coefficients were estimated by assigning 5500, 1490 and 125 $M^{-1}cm^{-1}$ units to each tryptophan, tyrosine, and cysteine residue, respectively, and summing those values. Peptides with a conjugated tetramethylrhodamine were quantified by absorption at 555 nm with an estimated molar extinction coefficient of 90,000 $M^{-1}cm^{-1}$. Peptides were then aliquoted into 1.5 mL polypropylene centrifuge tubes and lyophilized. Peptide aliquots were dissolved in DMSO to a concentration of 1 mM for later use and stored frozen with refrigeration.

#### C. Peptide Characterization:

Peptide 1

10-74% MeCN over 6 min. 0.3 mL/min

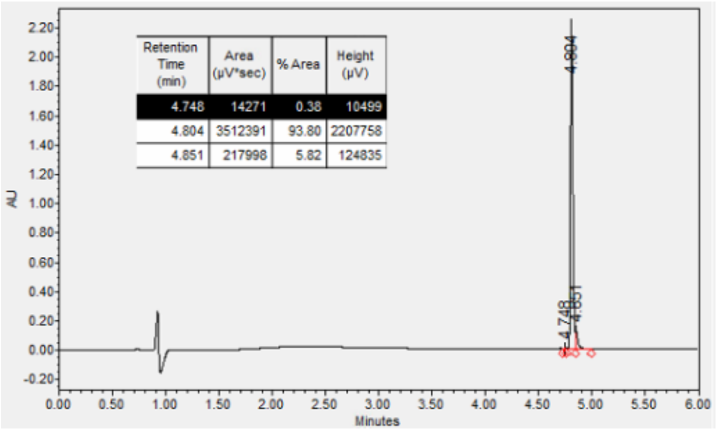

UPLC: 10-74% MeCN over 6 min. 0.3 mL/min, 93.8%

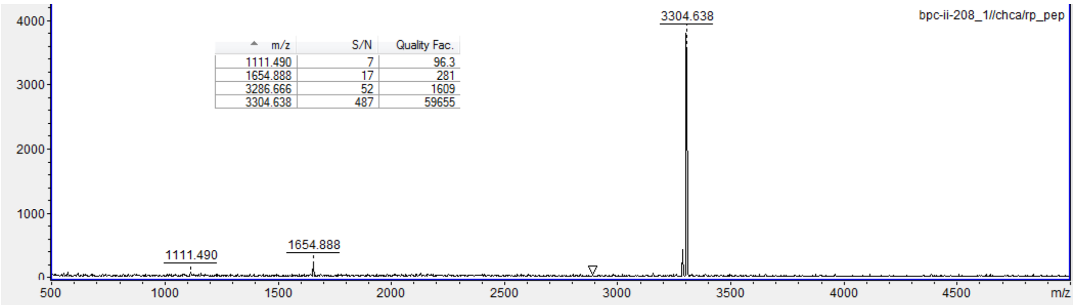

MALDI-TOF MS: For M = $C_{151}H_{228}N_{40}O_{44}$, Calc. $[M+H]^{+1}$ monoisotopic exact mass (*m*/*z*): 3306.69, found 3304.64

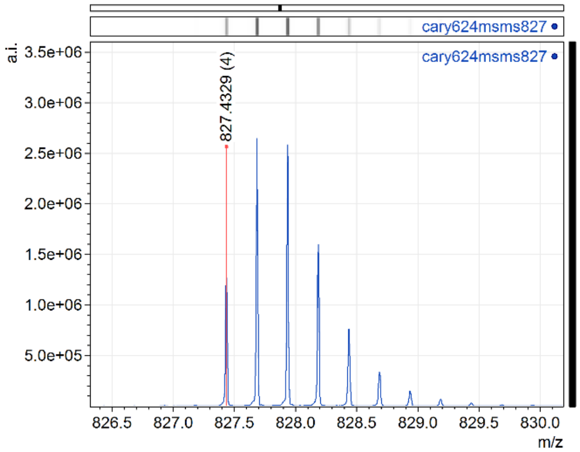

ESI-qTOF MS: Calc. monoisotopic $[M+4H]^{+4}$ (*m*/*z*): 827.4281, Found 827.4329

Peptide 2

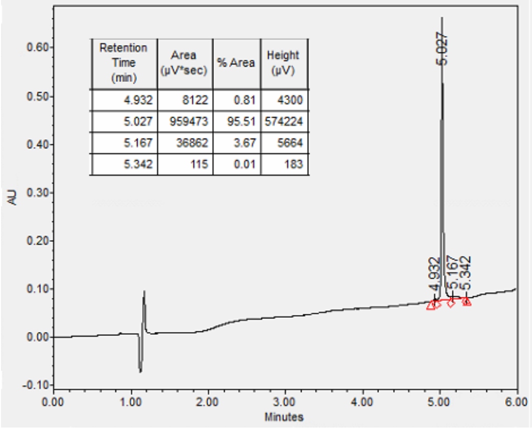

UPLC: 10-74% MeCN over 6 min. 0.3 mL/min, 95.5%

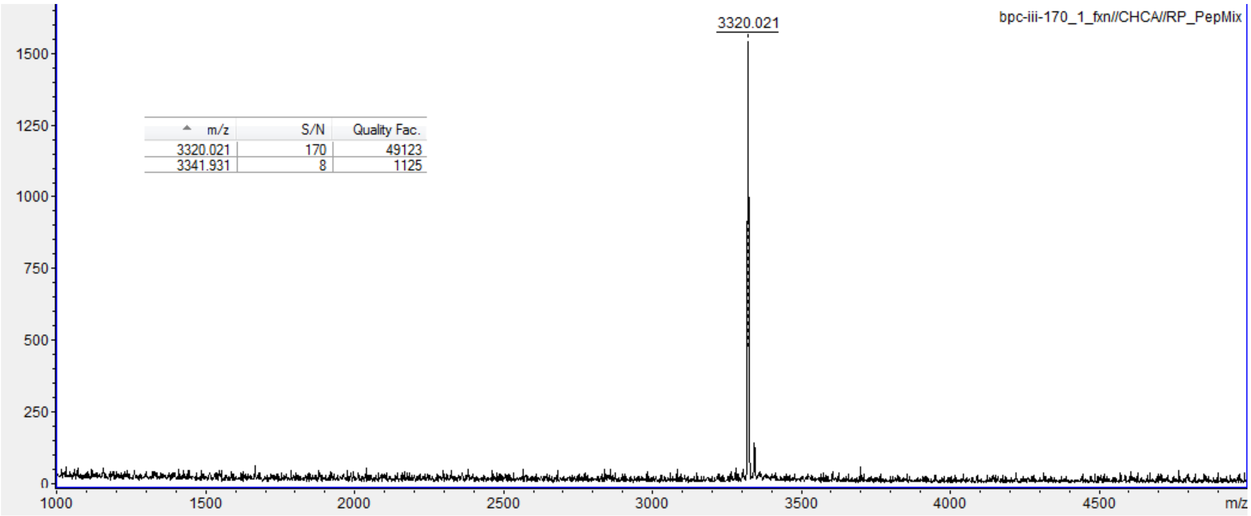

For M = $C_{152}H_{230}N_{40}O_{44}$, calc $[M+H]^{+1}$ (*m*/*z*): 3320.70, found 3320.0

Peptide 3

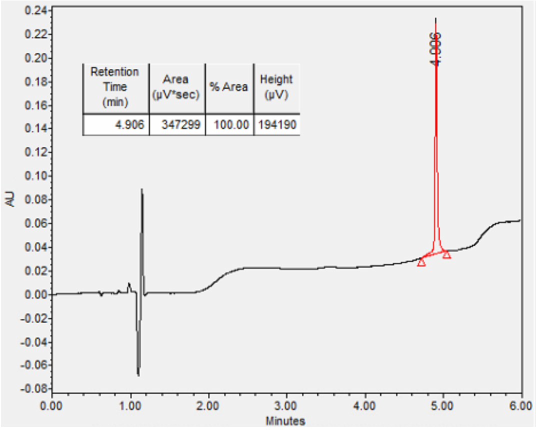

UPLC: 10-74% MeCN over 6 min. 0.3 mL/min, >99%

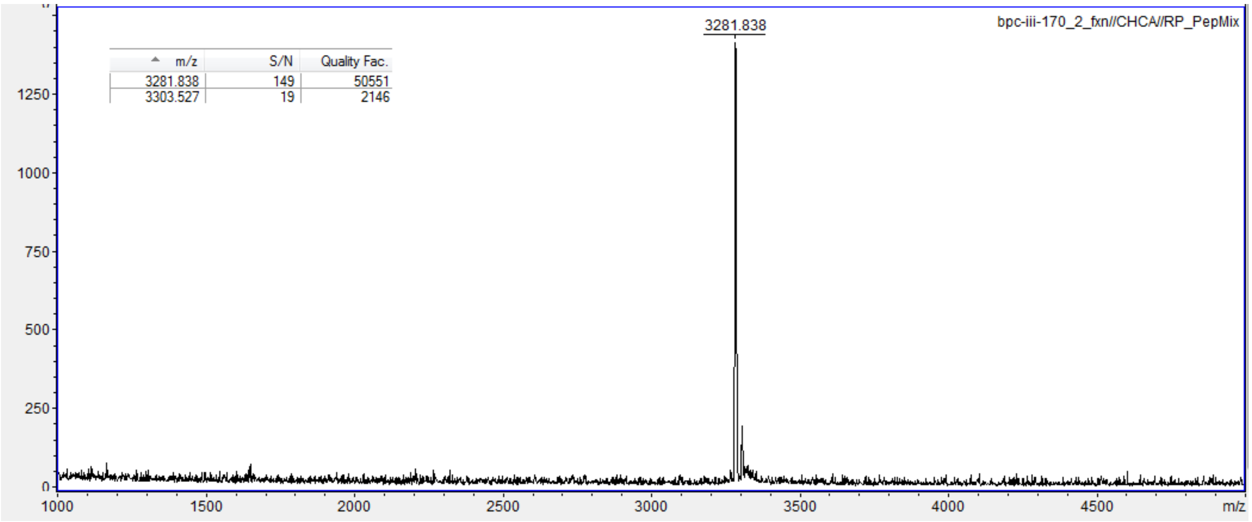

For M = $C_{148}H_{224}N_{40}O_{45}$, Calc. $[M+H]^{+1}$ monoisotopic exact mass (*m*/*z*): 3282.65, found 3281.83

Peptide 4

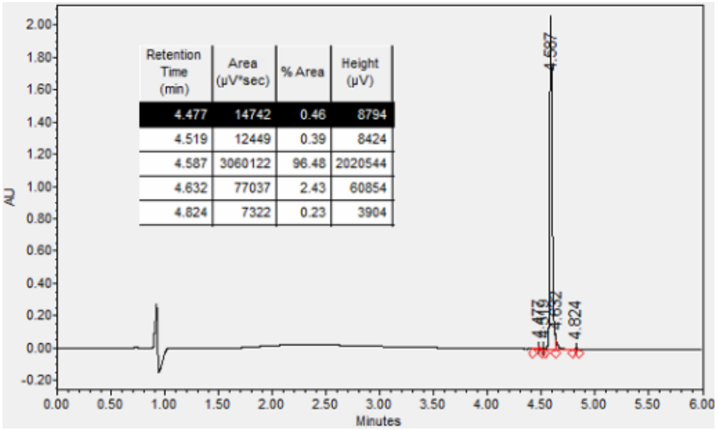

UPLC: 10-74% MeCN over 6 min. 0.3 mL/min, 96.5%

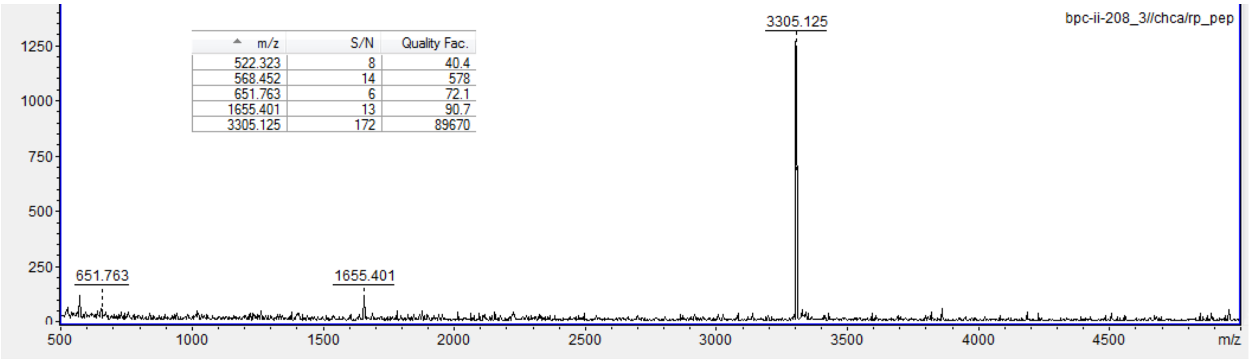

MALDI-TOF MS: For M = $C_{150}H_{227}N_{41}O_{44}$, Calc. $[M+H]^{+1}$ monoisotopic exact mass (*m/z*): 3307.69, found 3305.12

Peptide 5

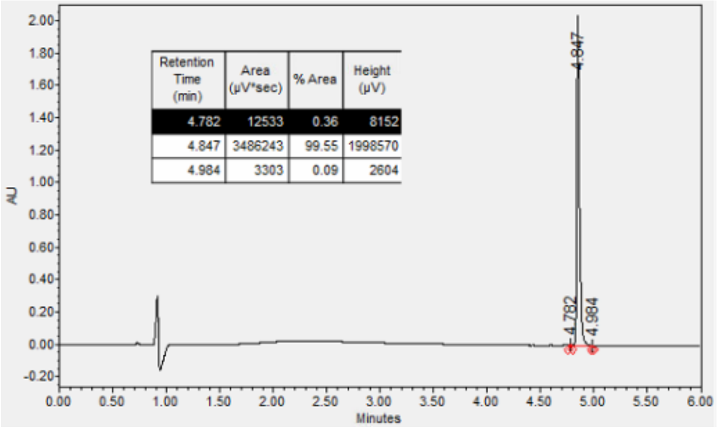

UPLC: 10-74% MeCN over 6 min. 0.3 mL/min, >99%

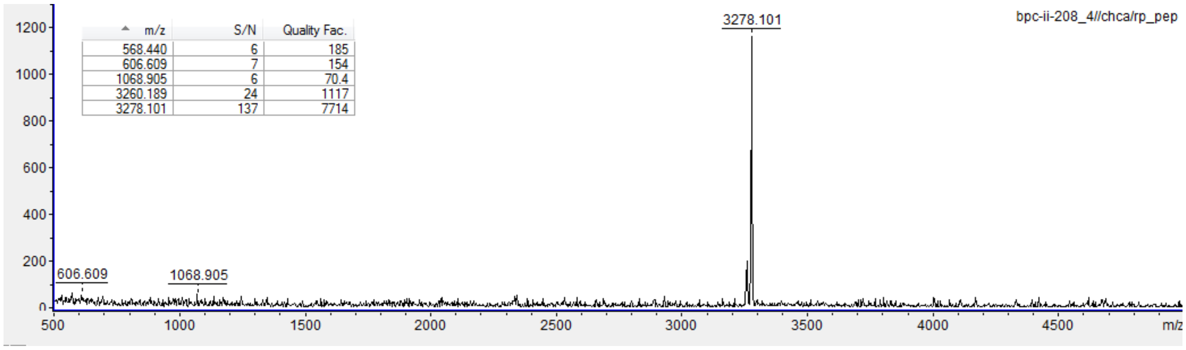

MALDI-TOF MS: For M = $C_{149}H_{226}N_{40}O^{44}$, Calc. $[M+H]^{+1}$ monoisotopic exact mass (*m*/*z*): 3280.67, found 3278.10

Peptide 6

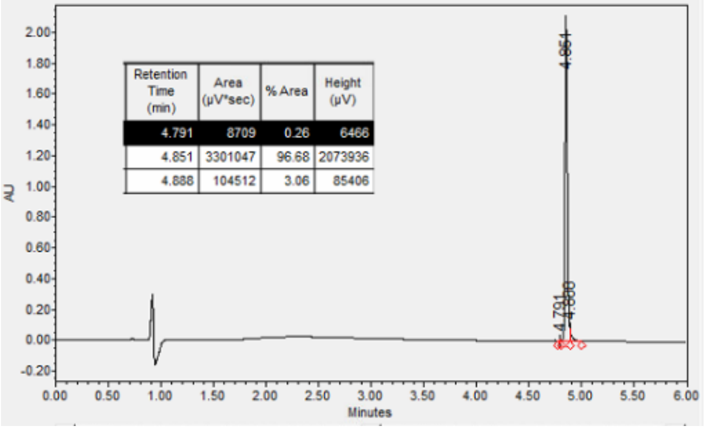

UPLC: 10-74% MeCN over 6 min. 0.3 mL/min, 96.7%

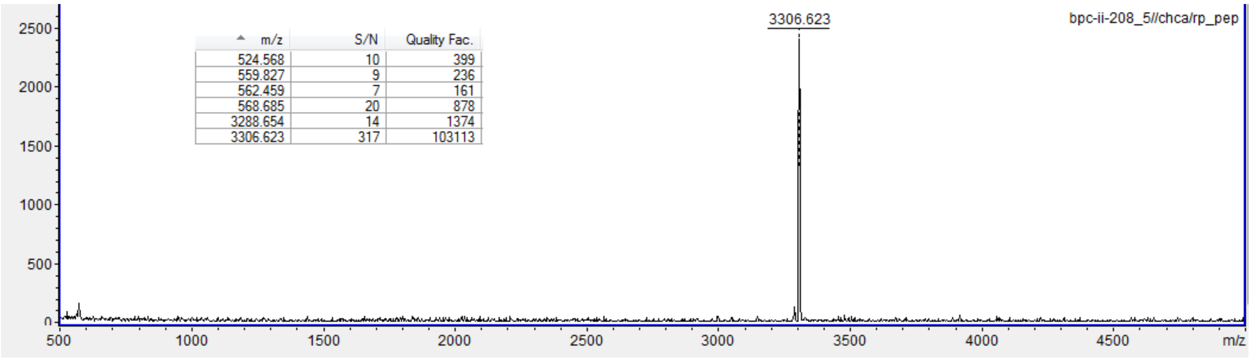

MALDI-TOF MS: For M = $C_{151}H_{230}N_{40}O_{44}$, Calc. $[M+H]^{+1}$ monoisotopic exact mass (*m*/*z*): 3308.71, found 3306.6.

Peptide 7

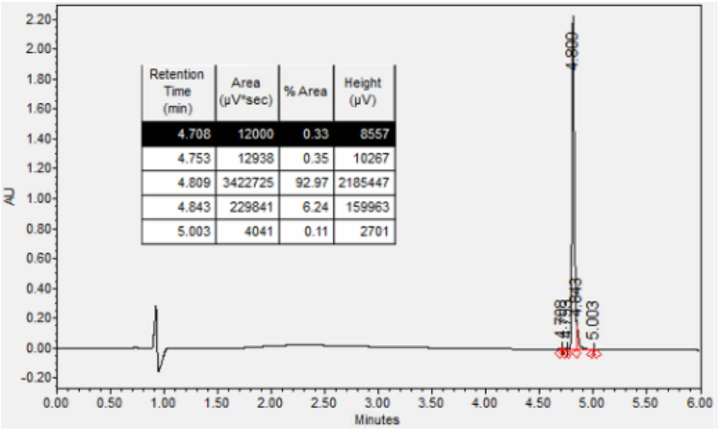

UPLC: 10-74% MeCN over 6 min. 0.3 mL/min, 92.9%

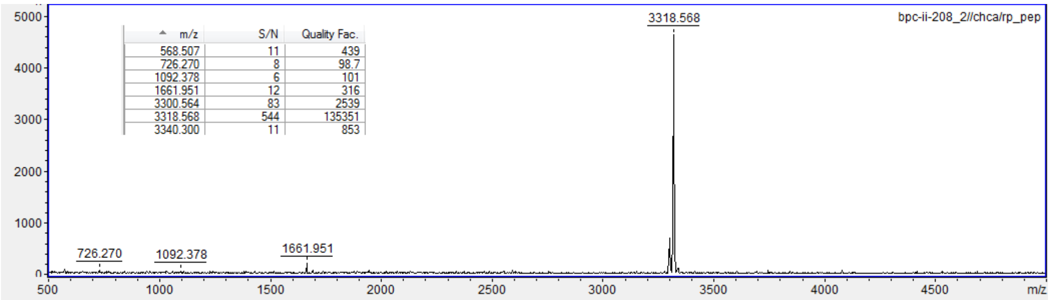

MALDI-TOF MS: For M = $C_{152}H_{230}N_{40}O_{44}$, Calc. $[M+H]^{+1}$ monoisotopic exact mass (*m*/*z*): 3320.71, found 3318.57

GLP-1-TMR:

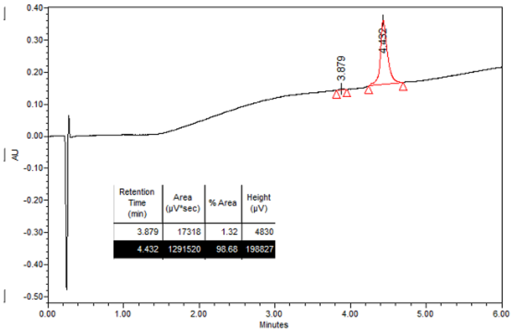

LC/MS: 10-90% MeCN over 6 min. 99%

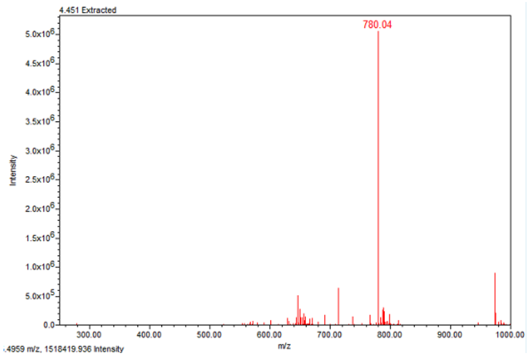

LC/MS, ESI-single quadrupole: For M = $C_{182}H_{261}N_{45}O_{51}$, Calc. $[M+5H]^{+5}$ most abundant *m/z* 779.79, found 780.04.

1-TMR:

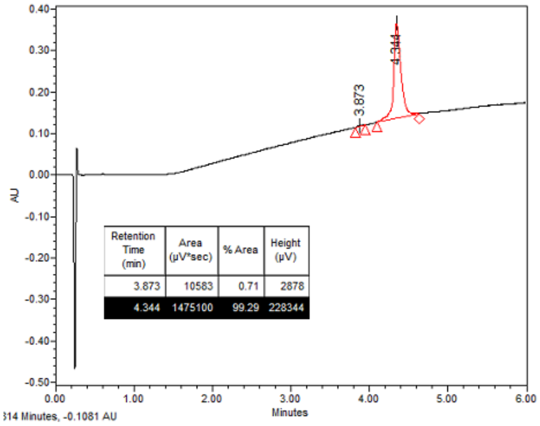

LC/MS: 10-90% MeCN over 6 min >99%

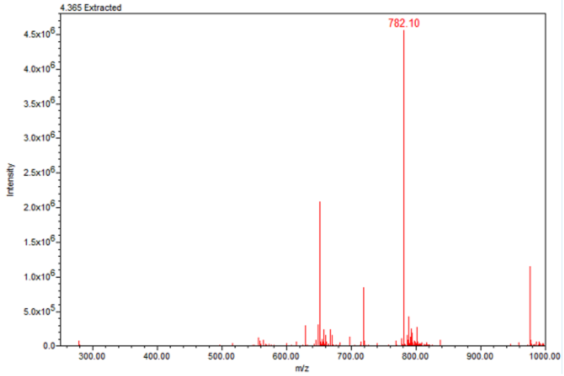

LC/MS, ESI-single quadrupole: For M = $C_{184}H_{263}N_{45}O_{50}$, Calc. $[M+5H]^{+5}$ most abundant *m/z* 781.80, found 782.10
